# Stochastic Variational Inference for Bayesian Phylogenetics: A Case of CAT Model

**DOI:** 10.1101/358747

**Authors:** Tung Dang, Hirohisa Kishino

## Abstract

The pattern of molecular evolution varies among gene sites and genes in a genome. By taking into account the complex heterogeneity of evolutionary processes among sites in a genome, Bayesian infinite mixture models of genomic evolution enable robust phylogenetic inference. With large modern data sets, however, the computational burden of Markov chain Monte Carlo sampling techniques becomes prohibitive. Here, we have developed a variational Bayesian procedure to speed up the widely used PhyloBayes MPI program, which deals with the heterogeneity of amino acid profiles. Rather than sampling from the posterior distribution, the procedure approximates the (unknown) posterior distribution using a manageable distribution called the variational distribution. The parameters in the variational distribution are estimated by minimizing Kullback-Leibler divergence. To examine performance, we analyzed three empirical data sets consisting of mitochondrial, plastid-encoded, and nuclear proteins. Our variational method accurately approximated the Bayesian phylogenetic tree, mixture proportions, and the amino acid propensity of each component of the mixture while using orders of magnitude less computational time.

## 1 Introduction

Understanding the evolutionary variation of phenotypic characters and testing hypotheses about the underlying mechanism are some of the main concerns of evolutionary biology. Because this variation needs to be interpreted as an evolutionary history, accurately inferring the phylogenetic tree is important. Otherwise, the uncertainty of phylogenetic inference must be taken into account to obtain an unbiased picture of evolutionary variation.

The increasing amount of available genomic data enables reliable inference of phylogenetic trees. Because molecular evolution is largely driven by nearly neutral or slightly deleterious mutations Ohta (1973), this process is less prone to convergent evolution than the evolution of phenotypic traits. The pattern of molecular evolution is statistically formulated by Markov processes. The pattern and rate of molecular evolution are complex, however, depending on various factors affecting mutation rates and functional constraints. To model protein evolution, Thorne, Goldman, and Jones (1996) introduced the concept of hidden states of secondary structure to describe sites of heterogeneity Goldman *et al.* (1996); Thorne *et al.* (1996); Jones *et al.* (1996). Koshi and Goldstein (1998) developed a model of the physico-chemical properties of amino acids, while Halpern and Bruno (1998) introduced a more advanced model with position-specific amino acid frequencies.

Equilibrium amino acid frequencies, which reflect structural and functional constraints, vary among sites within and among proteins. Inter-species comparative genomics approaches can analyze a huge number of alignment columns, but the number of taxa is often insufficient to estimate individual position-specific amino acid frequencies. To achieve a balance between variance and bias, Lartillot and Philippe (2004) proposed a Bayesian non-parametric approach based on a countable infinite mixture model, referred to as the CAT model. This model specifies K distinct processes (or classes), each characterized by a particular set of equilibrium frequencies, and sites are distributed according to a mixture of these K distinct processes. By proposing a truncated stick-breaking representation of the Dirichlet process prior on the space of equilibrium frequencies (Ferguson 1973; Green and Richardson 2001; Ishwaran and James 2001), the total number of classes can be treated as free variables of the model. A hybrid framework combining Gibbs-sampling and the Metropolis-Hastings algorithm has been developed to estimate all parameters of the model Papaspiliopoulos and Roberts (2008).

Existing approaches cannot take full advantage of the CAT model (Lartillot and Philippe 2004; Lartillot 2006), because the computational burden is prohibitive for inference based on large data sets. Even well-designed sampling schemes need to generate a large number of posterior samples through the entire data set to resolve convergence, and their convergence can be difficult to diagnose. To provide faster estimation, Lartillot *et al.* (2013) developed a message passing interface (MPI) for parallelization of the PhyloBayes MPI program. By implementing Markov chain Monte Carlo (MCMC) samplers in a parallel environment, PhyloBayes MPI allows for faster phylogenetic reconstruction under complex mixture models.

## 2 New Approaches

Here, we propose an alternative approach, a variational inference method (Jordan *et al.* 1999; Bishop 2006; Blei *et al.* 2006; Hoffman *et al.* 2013). Variational methods, originally used in statistical physics to approximate intractable integrals, have been successfully used in a wide variety of applications related to complex networks (Gopalan and Blei 2013) and population genetics (Gopalan *et al.* 2016; Raj *et al.* 2014). The basic idea of variational inference in the Bayesian framework is to approximate the posterior distribution by a computationally tractable function, called the variational distribution. The variational parameter, which specifies the variational distribution, is estimated by minimizing the Kullback-Leibler (KL) divergence of the posterior distribution to the variational distribution. As a result, the posterior distribution is estimated by numerical optimization without provoking Monte Carlo simulation. To deal with the uncertainty of tree topologies, we preserved the Gibbs sampling algorithm of tree topologies (Lartillot *et al.* 2013). In this article, we demonstrate that our algorithms are considerably faster than PhyloBayes MPI while achieving comparable accuracy.

### 2.1 Variational Inference of CAT-Poisson Model

In the CAT model, each site category has its own amino acid replacement rate matrix. Instead of dealing with the general time reversible Markov process, in this paper, we focus on the most popular CAT-Poisson model. This model takes account of rate heterogeneity among sites, and also allows the preferred amino acids to vary among sites. It assigns the alignment columns to the categories of amino acid profiles, taking account of uncertainty. Given the assignment to the category, the process of molecular evolution follows the amino acid version of the F81 model (Felsenstein 1981).

We denote the sequence data set by *D*. The CAT model has parameters (Φ, Ξ). Φ consists of branch lengths (*l*), site-specific relative rates (*r*), the amino acid profile (equilibrium frequency, *π*), the unit length of the stick (*V*) and the allocation variable (*z*) of the Dirichlet process prior on these profiles. The parameter Ξ is the substitution mapping parameter. Variational inference approximates the true intractable posterior distribution *p*(Φ, Ξ| *D*) by an element of a tractable family of probability distributions *q*(Φ, Ξ|Θ), called the *variational distribution*. As a variational distribution for the CAT-Poisson model, we adopt Gamma distributions for the branch lengths and the site-specific evolutionary rates, and a Dirichlet distribution for the amino acid profiles (see Materials and Methods for detail).

The distribution is parameterized by free parameters, called *variational parameters* Θ. Variational inference fits these parameters to find a distribution close to the true intractable posterior distribution of interest. The distance between the distributions *q*(Φ, Ξ|Θ) and *p*(Φ, Ξ|*D*) is measured by Kullback-Leibler (KL) divergence:

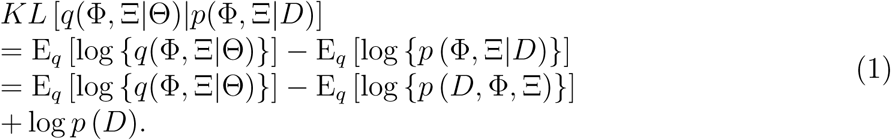

The term log *p* (*D*) in equation (1), which is the cause of computational difficulty in Bayesian analysis, can be treated as a constant term in numerical optimization to estimate the variational parameter:

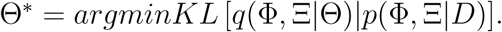

The variational inference maximizes the computational feasible target function:

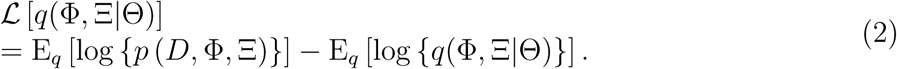

Because log *p* (*D*) *<* 0 and

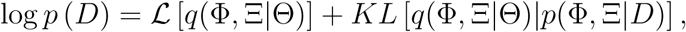

The equation (2) is called Evidence Lower BOund (ELBO; Jordan *et al.* (1999)).

It should be noted that, in the likelihood framework, a maximum likelihood approach minimizes the Kullback-Leibler divergence from the true distribution to the model distribution (Kullback and Leibler 1951; Akaike 1974). In contrast, a variational inference minimizes the Kullback-Leibler divergence from the model variational distribution to the true posterior distribution. Because of asymmetry of Kullback-Leibler divergence, the maximum value of ELBO cannot be used for comparing candidate models of variational distributions. Currently, the standard model checking process is to compare the important aspects of *q\**(Φ, Ξ|Θ) with those of MCMC runs by example data at the developmental stage of the program.

### 2.2 An Illustrative Example in Phylogenetics

As an illustrative example, we estimate the posterior distribution of the distance *d* between a pair of aligned sequences D with the JC69 model (Jukes and Cantor 1969). Out of *n* sites, the sequences differ at *x* sites. We assign independent gamma priors with *α* = 1 and *β* = 1 for the distance *d*:

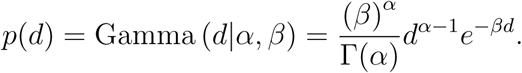

The likelihood of the JC69 model is given as:

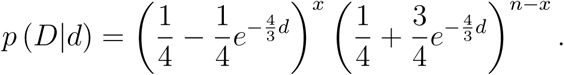

Given the prior and the likelihood, the posterior distribution is obtained as

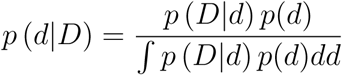

Because this illustrative model includes only a single free parameter, the denominator can be accurately calculated by numerical integration.

As the variational distribution for the posterior distribution of *d*, we adopt a gamma distribution:

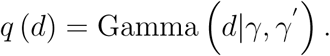

The ELBO is written as

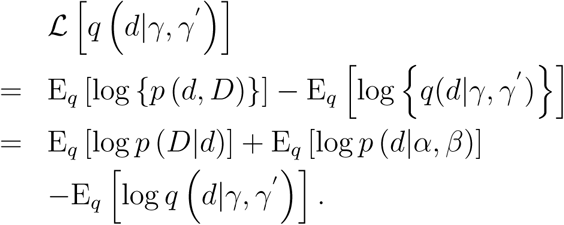

Therefore, variational parameters, *γ* and *γ’*, are estimated by optimizing the value of the following:

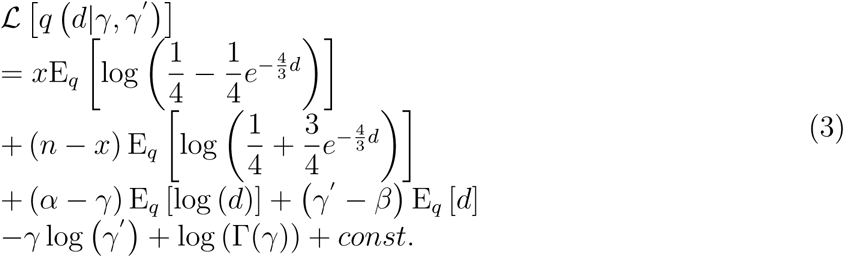

Here,

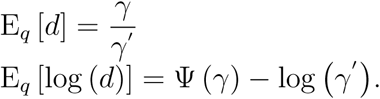

Ψ(.) is the digamma function, the first derivative of the log gamma function. The first term and the second term of eq. (3) are calculated by numerical integration. The variational parameters *γ* and *γ’* are estimated by maximizing eq.(3) numerically. For complex models with a large number of parameters, mathematical expansions such as the Taylor expansion (Ma and Leijon 2011; Ma *et al.* 2014) and the Delta method (Braun and McAuliffe 2010; Wang and Blei 2013) are often applied to integrands so that explicit forms of expectations are available.

Figure 1 shows the estimated posterior distribution of *d* for the case of *n* = 1000, *x* = 100, *α* = 1, *β* = 1. The distribution with the estimated parameters 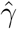 and 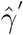 approximates the true posterior distribution accurately.

**Figure 1:**
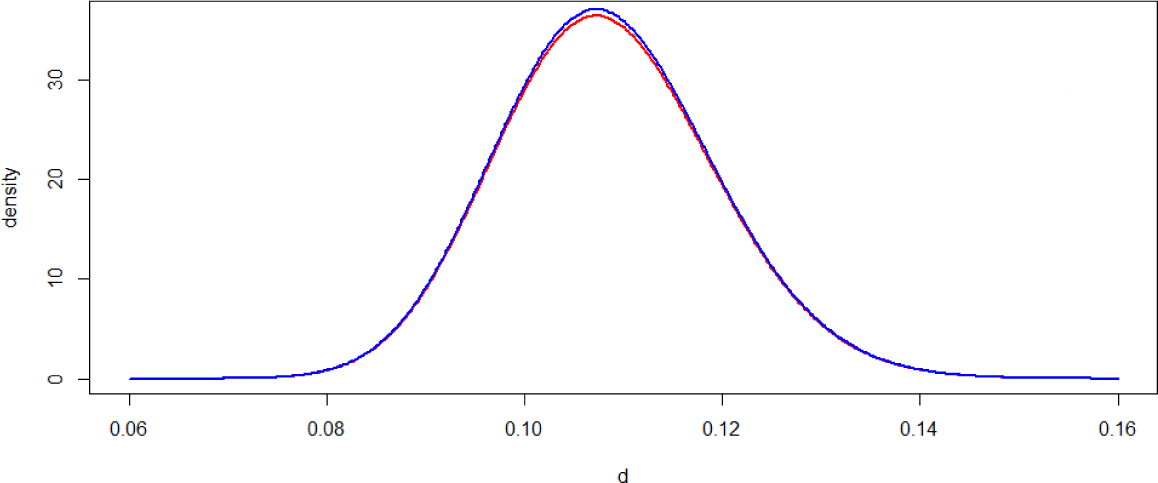
The variational inference of the posterior distribution of distance. The red curve is the estimated posterior distribution by variation inference, and the blue curve is the true posterior distribution.

## 3 Results

### 3.1 Runtime Performance

Table 1 compares the computational time of variational inference of the CAT-Poisson model with that of MCMC. Three empirical data sets were analyzed (Materials and Methods). Here, the number of iterations was set to 30,000 for MCMC sampling from the posterior distribution (default value of phyloBayes). As for variational inference, we could not implement a stopping rule based on convergence criteria because we partially preserved MCMC for tree topology. The trace of ELBO value implied sufficient convergence with far less than 1000 iterations for data set A (Figure 2(a)). However, we note that the value of ELBO expresses the goodness of fit of the variational parameters, but does not measure the consistency of the topology. Figures 2(b-d) show that the posterior consensus tree by variational inference mostly reached convergence at 1000. Tentatively, we set the same number of iterations as an MCMC case for comparing CPU times. We confirmed that the result of variational inference with 5000 iterations was unchanged for data set A (data not shown).

**Table 1:** Run times and estimated trees of variational inference and MCMC algorithms on real data

| Data set | Taxa | Sites | States | MCMC | VI |
| --- | --- | --- | --- | --- | --- |
| Data Set A | 13 | 6,622 | 20 | 4.72 days | 0.81 day |
| Data Set B | 28 | 10,137 | 20 | 10.61 days | 2.36 days |
| Data Set C | 66 | 38,330 | 20 | 28.35 days | 5.67 days |

Both variational inference and MCMC algorithms were run in a parallel environment. The properties of the parallel version were evaluated on a personal computer (Intel Core i7-6700 CPU 3.40GHz, 8 cores, 2 threads per core, 4 cores per socket, 16 Gb RAM), under Linux Mint 17.3 Rosa. In this comparative study, both variational inference and MCMC had 30,000 iterations (see text).

**Figure 2:**
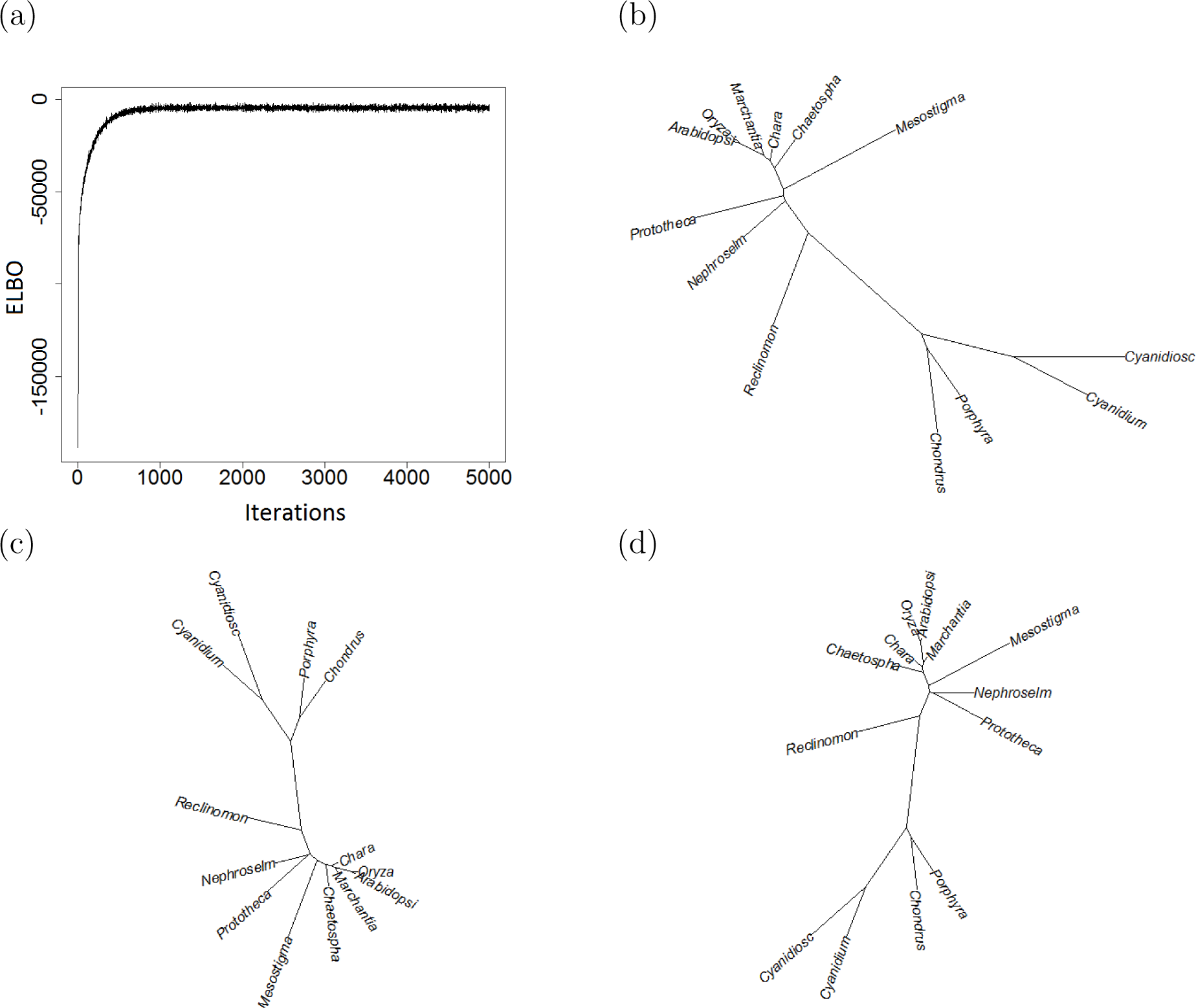
Convergence of variational inference for the mitochondrial data set (13 taxa and 6622 amino acid positions (Rodríguez-Ezpeleta *et al.* 2006)). The figures show the trace of ELBO value (a), and the estimated posterior consensus trees with 1000 iterations (b) and with 5000 iterations (c), in contrast to the result of 30000 iterations of MCMC (d).

The time complexity of each of the above algorithms was found to increase regularly with the numbers of genes, species, and total aligned amino acid positions. Even with the same number of iterations, run times were significantly reduced in the variational inference framework compared with those in the MCMC approach. This may be partly because variational inference does not include the step of generating random numbers (except for the one for sampling topologies) and the calculation of acceptance probabilities. Since our stopping rule was not thoughtfully designed but rather ad hoc, we need to perform any interpretations with caution. Once we can replace the step of Gibbs sampling of topology with some deterministic procedure of variational inference, the computational burden will be markedly reduced.

### 3.2 Posterior Independence between the Phylogenetic Parameters

Our variational distribution for the CAT model assumed independence among the branch lengths, the site-specific relative rates, and the amino acid profiles. To examine its validity, we checked the MCMC sample of the total branch length and the entropy of the amino acid profile of the largest cluster as an example. The scatter plot supports independence between these two characters (*r* = –0.024, Figure 3). As a result, the variational inference approximated the distribution of the MCMC sample accurately (Figure 4a,b). (The good fitting for each branch length can be seen in Figure S1.)

**Figure 3:**
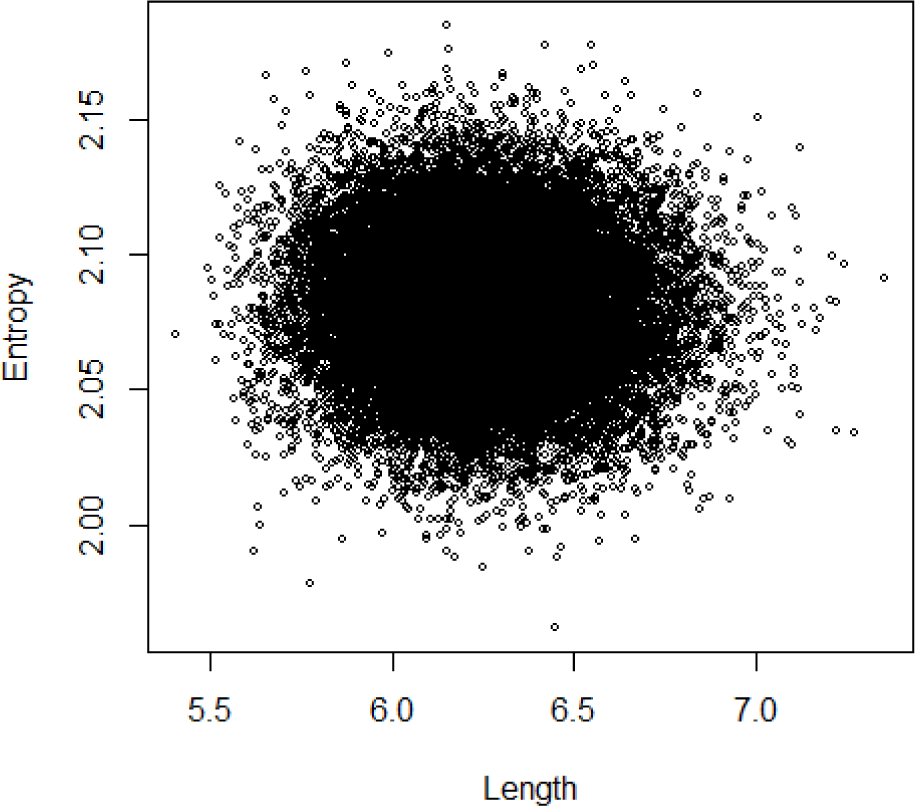
MCMC joint distributions of the total branch length and the entropy of the amino acid profile of the largest cluster based on the mitochondrial data set (13 taxa and 6622 amino acid positions (Rodríguez-Ezpeleta *et al.* 2006)).

**Figure 4:**
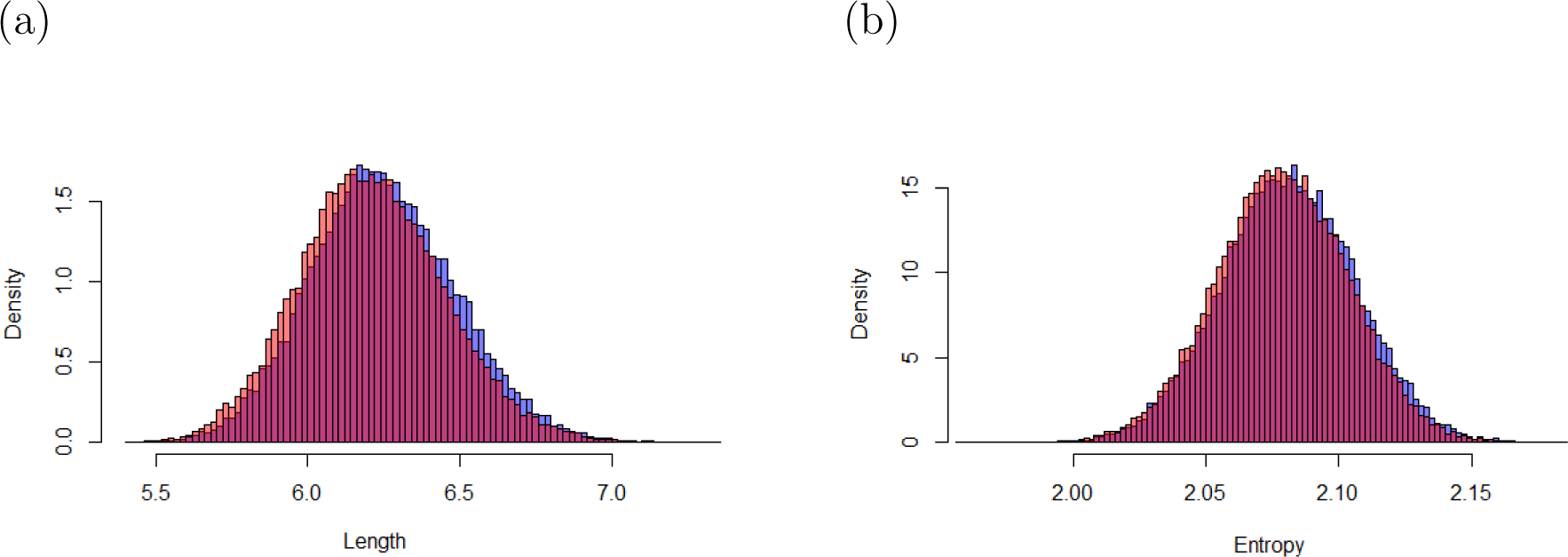
MCMC-based and variational inference-based posterior distributions of the total branch length and the entropy of the amino acid profile of the largest cluster based on the mitochondrial data set (13 taxa and 6622 amino acid positions (Rodríguez-Ezpeleta *et al.* 2006)). (a) The total branch length and (b) the entropy of the amino-acid profile of the largest category. Blue and red histograms are the distributions of the samples by MCMC and by variational inference, respectively.

### 3.3 Accuracy of Estimated Profiles

By introducing a Dirichlet process prior, the CAT model provides a posterior distribution of *K*, the number of separate categories, and the size of each category. The PhyloBayes MPI program, which is based on a hybrid strategy combining Gibbs sampling and Metropolis-Hastings algorithm, first proposes allocation variables and amino-acid profiles. The site to category allocation are sampled with the posterior weights of the mixture and profiles associated with each component of the mixture. Metropolis-Hastings algorithms are then used to sample the classes for sites. In contrast, our variational inference estimates the posterior distributions of the allocation variable for each site, weight, and amino acid profile of the categories.

Table 2 compares some major categories estimated by MCMC and variational inference. The size of each category was approximated by the number of sites assigned to that class. The number of distinct categories was estimated for data set A representing 6622 amino acid positions. As can be seen in the table, variational inference accurately approximated the posterior means of these category sizes. The posterior distributions of the number of site categories and the amino acid profiles are also well approximated by the variational inference (Figure 5).

**Table 2:** The size (number of sites) of large categories estimated by variational inference and MCMC in data set A

| Category | MCMC | VI | Category | MCMC | VI |
| --- | --- | --- | --- | --- | --- |
| 1 | 524 | 527 | 9 | 256 | 246 |
| 2 | 481 | 480 | 10 | 235 | 240 |
| 3 | 457 | 454 | 11 | 226 | 220 |
| 4 | 403 | 400 | 12 | 197 | 188 |
| 5 | 328 | 326 | 13 | 161 | 157 |
| 6 | 284 | 290 | 14 | 148 | 145 |
| 7 | 273 | 276 | 15 | 140 | 137 |
| 8 | 265 | 275 | 16 | 138 | 126 |

Top-ranked estimated categories are listed along with the number of sites distributed in each class. The results are for real data set A, with the number of sites calculated by counting sites allocated to each category.

**Figure 5:**
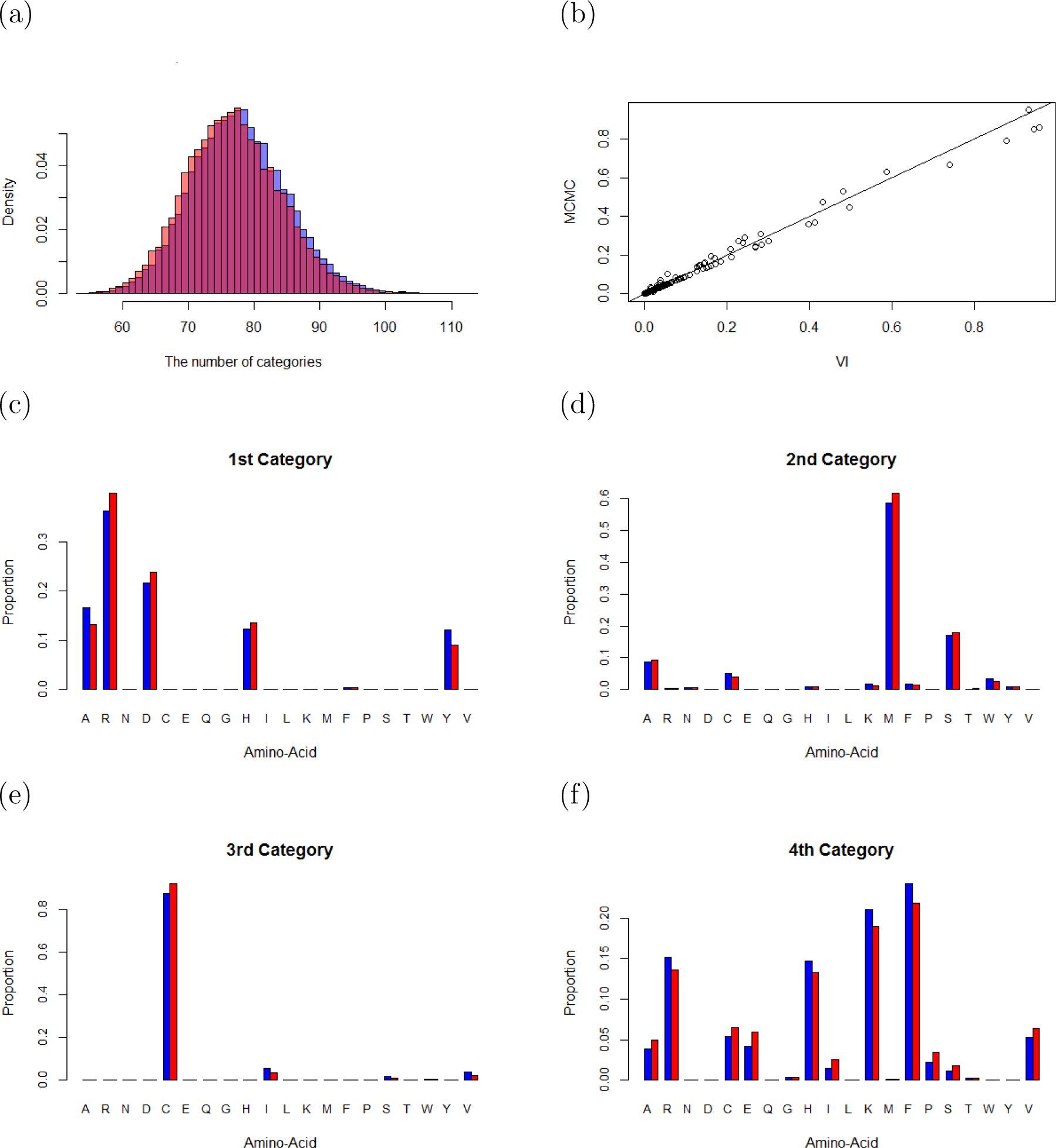
The MCMC and variational inference-based posterior distributions of the number of site categories and the amino acid profiles based on a mitochondrial data set (13 taxa and 6622 amino acid positions (Rodríguez-Ezpeleta *et al.* 2006)). (a) The posterior distributions of the number of site categories, (b) mean amino acid profiles of the first 16 site categories in Table 2 by variational inference vs. MCMC (c-f) posterior mean amino acid profiles of the four main site categories. Blue and red colors are the posterior means by MCMC and by variational inference, respectively.

Taken together, these results demonstrate that the estimation time required by the variational inference framework compares favorably with that used by sampling algorithms such as MCMC, while a sufficient level of accuracy under the CAT model is still guaranteed.

## 4 Discussion

The variational distribution for the CAT model approximated the posterior distribution accurately. This is largely because the branch lengths, site-specific evolutionary rates, and amino acid profiles were mostly independent in the posterior distribution. When the parameters of a model are mutually dependent in the joint posterior distribution, the variational inference may underestimate the posterior variance, even though the estimated posterior means may be unbiased. It is recommended to check the posterior correlations carefully at the stage of developing new programs, and to transform the parameters when the correlation is observed.

One of the most important steps in the variational framework is the calculations of expectations for the latent variables in the general ELBO. Specifically, the variational inference can achieve the best performances for the conjugate models. Because the likelihood of a CAT model is composed of the distributions of exponential family, most of the expectations could be obtained in the closed form.

The approximations of the posterior distributions of the transition probabilities in the Markov models of nucleotide substitution can still be a challenge for the Bayesian computation. There are some proposals that can deal with intractable integrations and provide a convenient way to obtain an analytically tractable solution, such as the first-order Taylor expansion (Ma and Leijon 2011; Ma *et al.* 2014) and the Delta method (Braun and McAuliffe 2010; Wang and Blei 2013), however, the mathematical expansions are still a challenge for the Bayesian phylogenetic inference. In many cases, phylogenetic inference includes many parameters, some of which are not of major concern. It may thus be worthwhile considering a practical approach to estimate these nuisance parameters by maximum likelihood and performing a Bayesian inference for the parameters of major interest.

## 5 Materials and Methods

### 5.1 CAT-Poisson Model

We briefly review the CAT-Poisson model that describes site heterogeneity of the substitution process (Lartillot and Philippe 2004). This model allows rate variation among sites and also allows variation of the rate matrix among sites. Here, we explain the basic default model, called the CAT-Poisson model. Given an amino acid sequence data set consisting of N alignment columns and P taxa, we denote the observed amino acid at site *i* for taxon *p* by *D*_*ip*_ (*i* = 1, *…, N*; 1 ≤ *p* ≤ *P*). The CAT-Poisson model regards the branch lengths *l*_*j*_ (1 < *j* < 2*P* − 3); the site-specific relative rates *r*_*i*_ (1 ≤ *i* ≤ *N*) as random variables. Each site has its specific amino acid profile, or equilibrium frequencies, *π*_*a*_, 1 ≤ *a* ≤ 20, such that 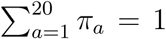. The subsitution process at each site follows the F81-type model (Felsenstein 1981). In other words, the probability of amino acid replacement by amino-acid *a* is proportional to *π*_*a*_. Sites are clustered into the categories of amino acid profiles. The CAT model describes the probabilistic allocation of a site to the categories by a mixture model. Given the allocation, the amino acid profile of a site has a prior of uniform Dirichlet distribution. A Dirichlet process treats the number of categories as an unknown variable. The stick-breaking representation considers two infinite collections of independent random variables; the unit length of sticks that correspond to the categories, *V*_*k*_, and the amino acid profiles of the categories, 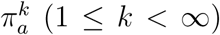. They follow:

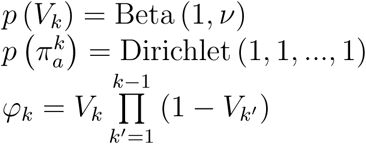

where *ϕ*_*k*_ is the mixing proportions of an infinite number of successively broken sticks and *v* stands for the total mass parameter of the Dirichlet process (Ferguson 1973; Green and Richardson 2001; Ishwaran and James 2001). Lartillot *et al*. (2013) introduced the allocation variable of a site *i* to a category, *z*_*i*_ *∈* [1,…, *∞*] (1 ≤ *i* ≤ *N*). The allocation variables are drawn i.i.d from a multinomial of the infinite vector of mixing proportions. Given that the site *i* belongs to the category *k*, the likelihood of the data at this site, *p*(*D*_*i*_|*π^k^*), is described by the transition probabilities along branches (Felsenstein 1981). *π^k^* is the amino acid profile of the *k*th category. Lartillot *et al*. (2013) applied a data augmentation algorithm of substitution mapping (Nielsen 2002). Along branch *j* and at site *i*, the substitution mapping, Ξ_*ij*_, is the combination of the number of substitutions, *n*_*ij*_, and the successive states of the process 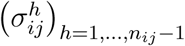. The random variable 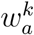 is the total number of substitutions to state *a* at sites that are assigned in category k, plus one if *a* is the state at the root of the tree. The prior distributions of the branch lengths and site-specific relative rates follow independent gamma distributions with shape 1 and scale *β* > 0 and independent gamma distributionswith shape *α* and scale *α*, respectively. *n*_*ij*_ follows the Poisson distribution with the rate parameter *r*_*i*_*l*_*j*_ and 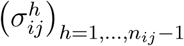 is drawn from 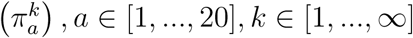.

### 5.2 Variational Inference of CAT Model

With mean-field variational approximations (Blei *et al*. 2006; Hoffman *et al*. 2013), each variable of the variational distribution is assumed to be independent. For practical implementation, we consider truncated stick-breaking representations (Blei *et al*. 2006) by setting the limit on the possible largest number of categories *K*_*max*_. The family of variational distributions in the CAT-Poisson model can be written as follows:

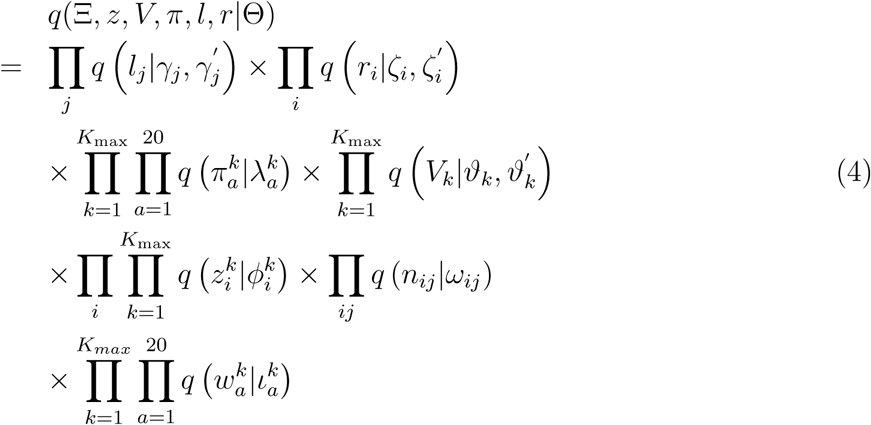

where

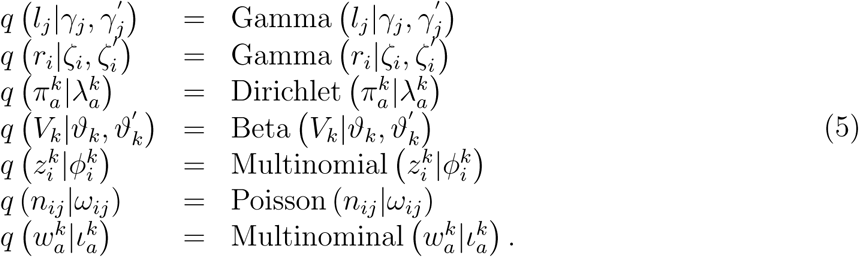

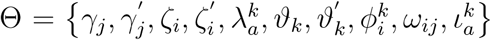 is the set of the free variational parameters. Note that eq (4) assumes independence among the sets of parameters describing phylogeny. This model may underestimate the posterior variance, if the true posterior joint distribution includes large correlations. We will see in the Result section that branch lengths, evolutionary rates, and amino acid profiles are almost independent in the joint distribution from MCMC. To guarantee the tractability of computing the expectations of variational distributions, we choose variational distributions from exponential families (Wainwright *et al*. 2008).

To estimate each variational parameter in the CAT-Poisson model (4,5), we consider dividing the set of variational variables into two subgroups - global variables [Φ_*g*_ = (Ξ, *π, l, r*)] and local variables [Φ_*l*_ = (*V, z*)]. The local variational variables (*V, z*) are per-data-point latent variables. The k^*th*^ local variable *V*_*k*_ is the unit length of k^*th*^ stick in the stick-breaking representation which is used to make the infinite vector of mixing proportions. The i^*th*^ local variable 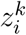 of the mixture component represents the allocation situation of site i of alignment of amino acid sequences. Each local variable 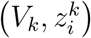 is governed by “local variational parameters” 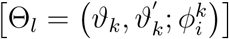. Bishop (2006) has proposed a coordinate ascent algorithm for solving the optimization problem of these variables. The coordinate ascent algorithm attempts to find the local optimum of the ELBO by optimizing each factor of the mean field variational distribution, while fixing the others. The optimal *q* (*z*) and *q* (*V)* are then proportional to the exponentiated expected log of the joint distribution,

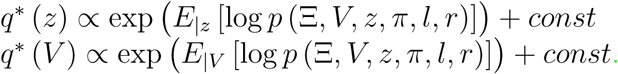

Here, *E*_|*z*_ and *E*_|*V*_ denote expectations with respect to the variational distributions of all the variables except for z or V. The global variables Φ_*g*_ potentially control any of the data. These variables are governed by the “global variational parameters” 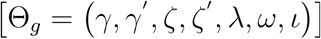. The coordinate ascent algorithm iterates t times to update local variational parameters based on mapping data,

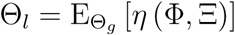

where *η* (.) are the natural parameters.

To estimate each global variational parameter in the CAT-Poisson model, we use the stochastic variational inference (SVI) algorithm to optimize the lower bound in Equation (2) (Hoffman *et al*. 2013). The stochastic variational algorithm is based on stochastic gradient ascent, the noisy realization of the gradient. In our study, we adopted natural gradients (Amari 1982) to account for the geometric structure of probability parameters (Robbins and Monro 1951). Importantly, natural gradients are easy to compute and give faster convergence than standard gradients. The SVI repeatedly subsamples the data, updates the values of the local parameters based on the subsampled data, and adjusts the global parameters in an appropriate way. Such estimates can guarantee algorithms to avoid shallow local optima of complex objective functions.

In our setting, we sample a mapping data point Ξ_*n*_ at each iteration, and compute the conditional natural parameters for the global variational parameters given N replicates of Ξ_*n*_. Then, the noisy natural gradients are obtained. By using these gradients, we update Θ_*g*_ at each of t iterations (with step size *ρ*_*t*_)

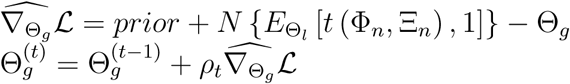

where *t* (.) denote the sufficient statistics.

Based on the subsampling techniques, this procedure reduces the computational burden by avoiding the expensive sums in the above lower bound. The SVI algorithm thus significantly accelerates the variational objective analysis of the large database. Applying the previously proposed SVI framework (Hoffman *et al*. 2013), we can separate the computational cycle into the following steps:

1. Sample amino acid data from the whole set of input data.
2. Estimate how each site is assigned to a category, based on observational data and the current approximation of variational parameters.
3. Update variational parameters
  - Local parameters are assignment variables, and breaking proportions.
  - Global parameters are equilibrium frequency profile, branch length, and rate across sites.

The lower bound of the data in terms of the variational parameters is specifically described in the Supplementary Material. Mathematical details of the variational objective function and computational methods of noisy derivatives and updating of variational parameters are also explained in that section.

### 5.3 Parallelization and Tree Topology

To parallelize the algorithm at the single machine level and thus reduce runtimes, we adopted the MPI parallelization of the PhyloBayes MPI program (Lartillot *et al*. 2013). Specifically, we used one master process for dispatching computational tasks and collecting and summing results, and with multiple slave processes executing the orders and returning all essential information to the master. This parallel strategy helps to equally divide the computational burden among slaves.

In addition, a partial Gibbs sampling algorithm for pruning and regrafting (SPR) is adopted to update the tree topology (Lartillot *et al*. 2013). In a parallel environment, the task of the master process is to randomly select a subtree for pruning and send this information to all slaves. The task of each slave process is to update the conditional likelihood vectors of each resulting topology and the complete scan of all possible regrafting points. One single log likelihood for each regrafting point is arranged into an array and sent back to the master process. All arrays are collected and summed and lastly the Gibbs sampling decision rule is finally applied to select the regrafting position.

### 5.4 Data Sets

Three real data sets were used for our computational experiments. Data set A was a mitochondrial data set consisting of 33 proteins and 6622 amino acid positions from 13 species. Data set B was a plastid data set composed of 50 plastidencoded proteins and 10137 amino acid positions from 28 species. In total, 13% and 5% amino acid positions were missing from the mitochondrial and plastid data sets, respectively (Rodriguez-Ezpeleta *et al*. 2006; Lartillot *et al*. 2013). Finally, data set C was a more challenging and larger complete set of mitochondrial protein sequences derived from a large alignment of EST and genome data, which consists of 197 genes and a total of 38330 amino acid positions from 66 species and with 30% missing data, constructed by (Philippe *et al*. 2011).

C++ code for the variational inference version of the CAT model to perform computational experiments with these data sets is available at https://github.com/tungtokyo1108/.

## Supporting information

Mathematical explanation

## 6 Acknowledgments

We thank the editor and two anonymous reviewers for constructive comments, all of which improved the manuscript significantly. We thank Edanz Group (www.edanzediting.com/ac) for editing the English text of a draft of this manuscript. This study was supported by Grantin-Aid for Scientific Research (B) 16H02788 from the Japan Society for the Promotion of Science.

## References

Akaike, H. 1974. A new look at the statistical model identification. IEEE Transactions on Automatic Control, 19: 716–723.

Amari, S.-I. 1982. Differential geometry of curved exponential families-curvatures and information loss. The Annals of Statistics, pages 357–385.

Bishop, C. M. 2006. Pattern recognition and machine learning. springer.

Blei, D. M., Jordan, M. I., et al. 2006. Variational inference for dirichlet process mixtures. Bayesian analysis, 1(1): 121–143.

Braun, M. and McAuliffe, J. 2010. Variational inference for large-scale models of discrete choice. Journal of the American Statistical Association, 105(489): 324–335.

Felsenstein, J. 1981. Evolutionary trees from dna sequences: a maximum likelihood approach. Journal of molecular evolution, 17(6): 368–376.

Ferguson, T. S. 1973. A bayesian analysis of some nonparametric problems. The annals of statistics, pages 209–230.

Goldman, N., Thorne, J. L., and Jones, D. T. 1996. Using evolutionary trees in protein secondary structure prediction and other comparative sequence analyses. Journal of molecular biology, 263(2): 196–208.

Gopalan, P., Hao, W., Blei, D. M., and Storey, J. D. 2016. Scaling probabilistic models of genetic variation to millions of humans. Nature genetics, 48(12): 1587.

Gopalan, P. K. and Blei, D. M. 2013. Efficient discovery of overlapping communities in massive networks. Proceedings of the National Academy of Sciences, 110(36): 14534–14539.

Green, P. J. and Richardson, S. 2001. Modelling heterogeneity with and without the dirichlet process. Scandinavian journal of statistics, 28(2): 355–375.

Halpern, A. L. and Bruno, W. J. 1998. Evolutionary distances for protein-coding sequences: modeling site-specific residue frequencies. Molecular biology and evolution, 15(7): 910–917.

Hoffman, M. D., Blei, D. M., Wang, C., and Paisley, J. 2013. Stochastic variational inference. The Journal of Machine Learning Research, 14(1): 1303–1347.

Ishwaran, H. and James, L. F. 2001. Gibbs sampling methods for stick-breaking priors. Journal of the American Statistical Association, 96(453): 161–173.

Jones, D. T., Orengo, C. A., and Thornton, J. M. 1996. Protein folds and their recognition from sequence. Protein structure prediction a practical approach, pages 173–204.

Jordan, M. I., Ghahramani, Z., Jaakkola, T. S., and Saul, L. K. 1999. An introduction to variational methods for graphical models. Machine learning, 37(2): 183–233.

Jukes, T. H. and Cantor, C. R. 1969. Evolution of protein molecules. In H. N. Munro, editor, Mammalian Protein Metabolism, pages 21–132. Academic Press.

Koshi, J. and Goldstein, R. 1998. Models of natural mutations including site heterogeneity. Proteins, 32(3): 289–295.

Kullback, S. and Leibler, R. A. 1951. On information and sufficiency. Annals of Mathematical Statistics, 22: 79–86.

Lartillot, N. 2006. Conjugate gibbs sampling for bayesian phylogenetic models. Journal of computational biology, 13(10): 1701–1722.

Lartillot, N. and Philippe, H. 2004. A bayesian mixture model for across-site heterogeneities in the amino-acid replacement process. Molecular biology and evolution, 21(6): 1095–1109.

Lartillot, N., Rodrigue, N., Stubbs, D., and Richer, J. 2013. Phylobayes mpi: phylogenetic reconstruction with infinite mixtures of profiles in a parallel environment. Systematic Biology, 62(4): 611–615.

Ma, Z. and Leijon, A. 2011. Bayesian estimation of beta mixture models with variational inference. IEEE Transactions on Pattern Analysis & Machine Intelligence, (11): 2160–2173.

Ma, Z., Rana, P. K., Taghia, J., Flierl, M., and Leijon, A. 2014. Bayesian estimation of dirichlet mixture model with variational inference. Pattern Recognition, 47(9): 3143–3157.

Nielsen, R. 2002. Mapping mutations on phylogenies. Systematic biology, 51(5): 729–739.

Ohta, T. 1973. Slightly deleterious mutant substitutions in evolution. Nature, 246(5428): 96.

Papaspiliopoulos, O. and Roberts, G. O. 2008. Retrospective markov chain monte carlo methods for dirichlet process hierarchical models. Biometrika, 95(1): 169–186.

Philippe, H., Brinkmann, H., Copley, R. R., Moroz, L. L., Nakano, H., Poustka, A. J., Wallberg, A., Peterson, K. J., and Telford, M. J. 2011. Acoelomorph flatworms are deuterostomes related to xenoturbella. Nature, 470(7333): 255.

Raj, A., Stephens, M., and Pritchard, J. K. 2014. faststructure: variational inference of population structure in large snp data sets. Genetics, 197(2): 573–589.

Robbins, H. and Monro, S. 1951. A stochastic approximation method. The annals of mathematical statistics, pages 400–407.

Rodríguez-Ezpeleta, N., Philippe, H., Brinkmann, H., Becker, B., and Melkonian, M. 2006. Phylogenetic analyses of nuclear, mitochondrial, and plastid multigene data sets support the placement of mesostigma in the streptophyta. Molecular Biology and Evolution, 24(3): 723–731.

Thorne, J. L., Goldman, N., and Jones, D. T. 1996. Combining protein evolution and secondary structure. Molecular Biology and Evolution, 13(5): 666–673.

Wainwright, M. J., Jordan, M. I., et al. 2008. Graphical models, exponential families, and variational inference. Foundations and TrendsQR in Machine Learning, 1(1–2): 1–305.

Wang, C. and Blei, D. M. 2013. Variational inference in nonconjugate models. Journal of Machine Learning Research, 14(Apr): 1005–1031.

