## Supplementary material for "Stochastic Variational Inference for Bayesian Phylogenetics: A Case of CAT Model": Mathematical explanation

*Tung Dang,*[*,1] *and Hirohisa Kishino*[1]
[1] Laboratory of Biometrics and Bioinformatics, University of Tokyo, Tokyo, Japan

#### A. Variational Inference for CAT model

#### 1 Mean-Field Variational Inference for CAT Model

We expand specifically the variational lower bound equation as follows:

$$\begin{array}{l}\mathcal{L} = \sum\limits_{k=1}^{K_{max}} \mathrm{E}_q\left[\log\left(p\left(V_k|1,\kappa\right)\right)\right] - \sum\limits_{k=1}^{K_{\max}} \mathrm{E}_q\left[\log\left(q\left(V_k|\vartheta_k,\vartheta_k^{'}\right)\right)\right]\\ +\sum\limits_i \sum\limits_{k=1}^{K_{max}} \mathrm{E}_q\left[\log p(Z_i^k|V_1,V_2,...,V_{K_{max}})\right] - \sum\limits_i \sum\limits_{k=1}^{K_{\max}} \mathrm{E}_q\left[\log\left(q\left(Z_i^k|\phi_i^k\right)\right)\right]\\ +\sum\limits_{k=1}^{K_{max}} \sum\limits_{a=1}^{20} \mathrm{E}_q\left[\log p(\pi_a^k|\upsilon_a)\right] - \sum\limits_{k=1}^{K_{\max}} \sum\limits_{a=1}^{20} \mathrm{E}_q\left[\log\left(q\left(\pi_a^k|\lambda_a^k\right)\right)\right]\\ +\sum\limits_j \mathrm{E}_q\left[\log\left(p\left(l_j|1,\beta\right)\right)\right] - \sum\limits_j \mathrm{E}_q\left[\log\left(q\left(l_j|\gamma_j,\gamma_j^{'}\right)\right)\right]\\ +\sum\limits_i \mathrm{E}_q\left[\log\left(p\left(r_i|\alpha,\alpha\right)\right)\right] - \sum\limits_i \mathrm{E}_q\left[\log\left(q\left(r_i|\zeta_i,\zeta_i^{'}\right)\right)\right]\\ +\mathrm{E}_q\left[\log\left(p\left(\Xi|V,z,\pi,l,r\right)\right)\right] - \mathrm{E}_q\left[\log\left(q\left(\Xi|\omega,\iota\right)\right)\right]\\ +\mathrm{E}_q\left[\log\left(p\left(D|\Xi,V,z,\pi,l,r\right)\right)\right]\end{array} \tag{1}$$

To compute the variational expectations $E_q[.]$ in equation (1), we use the properties of exponential family distribution. If variational distribution for branch lengths and rates $q\left(l_j|\gamma_j,\gamma_j^{'}\right)$, $q\left(r_i|\zeta_i,\zeta_i^{'}\right)$ are Gamma distributions, then the exponential family representations are given by

$$\begin{array}{l} q\left(l_j|\gamma_j,\gamma_j^{'}\right) = \exp\left(\gamma_j\log(\gamma_j^{'}) - \log\left(\Gamma(\gamma_j)\right) + (\gamma_j - 1)\log(l_j) - \gamma_j^{'} l_j\right)\\ q\left(r_i|\zeta_i,\zeta_i^{'}\right) = \exp\left(\zeta_i\log(\zeta_i^{'}) - \log\left(\Gamma(\zeta_i)\right) + (\zeta_i - 1)\log(r_i) - \zeta_i^{'} r_i\right)\end{array}$$

So the natural parameters of Gamma distribution for branch lengths and rates are $\eta_{l_j} = \left[\begin{array}{cc}\gamma_j - 1 & -\gamma_j^{'}\end{array}\right]$ and $\eta_{r_i} = \left[\begin{array}{cc}\zeta_i - 1 & -\zeta_i^{'}\end{array}\right]$. The sufficient statistics are $T\left(l_j\right) = \left[\begin{array}{cc}\log\left(l_j\right) & l_j\end{array}\right]$ and $T\left(r_i\right) = \left[\begin{array}{cc}\log\left(r_i\right) & r_i\end{array}\right]$. The approximation expectations are

$$\begin{array}{ll}\mathrm{E}_q\left[l_j\right] = \frac{\gamma_j}{\gamma_j^{'}} & \mathrm{E}_q\left[r_i\right] = \frac{\zeta_i}{\zeta_i^{'}}\\ \mathrm{E}_q\left[\log\left(l_j\right)\right] = \psi\left(\gamma_j\right) - \log\left(\gamma_j^{'}\right) & \mathrm{E}_q\left[\log\left(r_i\right)\right] = \psi\left(\zeta_i\right) - \log\left(\zeta_i^{'}\right)\end{array}$$

where $\psi(.)$ is the digamma function.
With equilibrium frequency profile, variational distributions $q\left(\pi_a^k|\lambda_a^k\right)$ follow the Dirichlet distributions, the exponential family representations are given by

$$q\left(\pi_k|\lambda_k\right) = \exp\left[\left(\sum_{a=1}^{20}\left(\lambda_a^k - 1\right)\log\left(\pi_a^k\right)\right) + \log\Gamma\left(\sum_{a=1}^{20}\lambda_a^k\right) - \sum_{a=1}^{20}\left(\log\Gamma\left(\lambda_a^k\right)\right)\right]$$

The natural parameters and sufficient statistics of Dirichlet distributions are $\eta_a^k = \lambda_a^k - 1$, $T\left(\pi_a^k\right) = \log\left(\pi_a^k\right)$. The approximation expectations are

$$\begin{array}{l}\mathrm{E}\left[\pi_a^k\right] = \frac{\lambda_a^k}{\sum\limits_{a'=1}^{20}\lambda_{a'}^k}\\ \mathrm{E}\left[\log\left(\pi_a^k\right)\right] = \psi\left(\lambda_a^k\right) - \psi\left(\sum\limits_{a'=1}^{20}\lambda_{a'}^k\right)\end{array}$$

With truncated stick-breaking representation of Dirichlet Mixture process, variational factors for the stick lengths $q\left(V_k|\vartheta_k, \vartheta'_k\right)$ are Beta distributions and assignment variable $z_i^k = \mathrm{I}\left[z_i = k\right]$ for the $\mathrm{i}^{th}$ site allocation is governed by a multinomial distribution indexed by a variational parameter $\phi_i^k$. The computation of the approximate expectation for parameters of truncated stick-breaking representation have been considered carefully by Blei and Jordan (2006). The results of variational expectation for both variables are as follows:

$$\begin{aligned}
&q\left(z_i = k\right) = \phi_i^k \\
&q\left(z_i > k\right) = \sum_{k'=k+1}^{K_{\max}} \phi_i^{k'} \\
&\mathrm{E}_q\left[\log\left(V_k\right)\right] = \psi\left(\vartheta_k\right) - \psi\left(\vartheta_k + \vartheta'_k\right) \\
&\mathrm{E}_q\left[\log\left(1 - V_k\right)\right] = \psi\left(\vartheta'_k\right) - \psi\left(\vartheta_k + \vartheta'_k\right)
\end{aligned}$$

Simplifying variational distributions for all parameters, we get

$$\begin{aligned}
\mathcal{L} = &\sum_{k=1}^{K_{\max}} \left\{\log\left[\Gamma\left(1+\kappa\right)\right] - \log\left[\Gamma\left(\kappa\right)\right] + \left(\kappa - 1\right)\left[\psi\left(\vartheta'_k\right) - \psi\left(\vartheta_k + \vartheta'_k\right)\right]\right\} \\
&- \sum_{k=1}^{K_{\max}} \left\{\begin{array}{l} \log\left[\Gamma\left(\vartheta_k + \vartheta'_k\right)\right] - \log\left[\Gamma\left(\vartheta_k\right)\right] - \log\left[\Gamma\left(\vartheta'_k\right)\right] + \left(\vartheta_k - 1\right)\left[\psi\left(\vartheta_k\right) - \psi\left(\vartheta_k + \vartheta'_k\right)\right] \\ + \left(\vartheta'_k - 1\right)\left[\psi\left(\vartheta'_k\right) - \psi\left(\vartheta_k + \vartheta'_k\right)\right] \end{array}\right\} \\
&+ \sum_{k=1}^{K_{\max}} \left\{\sum_i \phi_i^k \left(\psi\left(\vartheta_k\right) - \psi\left(\vartheta_k + \vartheta'_k\right)\right) + \left(\sum_{k'=k+1}^{K_{\max}} \left(\sum_i \phi_i^{k'}\right)\left(\psi\left(\vartheta'_{k'}\right) - \psi\left(\vartheta_{k'} + \vartheta'_{k'}\right)\right)\right)\right\} \\
&- \sum_{k=1}^{K_{\max}} \left\{\sum_i \phi_i^k \log\left(\phi_i^k\right)\right\} \\
&+ \left\{\left(\sum_{k=1}^{K_{\max}} \sum_a \left(\upsilon_a - 1\right)\left[\psi\left(\lambda_a^k\right) - \psi\left(\sum_{a'=1}^{20} \lambda_{a'}^k\right)\right]\right) + \log\Gamma\left(\sum_{a=1}^{20} \upsilon_a\right) - \textstyle\sum_{a=1}^{20}\left(\log\Gamma\left(\upsilon_a\right)\right)\right\} \\
&- \sum_{k=1}^{K_{\max}} \left\{\sum_{a=1}^{20} \left(\lambda_a^k - 1\right)\left[\psi\left(\lambda_a^k\right) - \psi\left(\sum_{a'=1}^{20} \lambda_{a'}^k\right)\right] + \log\Gamma\left(\sum_{a=1}^{20} \lambda_a^k\right) - \textstyle\sum_{a=1}^{20} \log\Gamma\left(\lambda_a^k\right)\right\} \\
&+ \sum_j \left\{\log(\beta) - \beta\left[\frac{\gamma_j}{\gamma'_j}\right]\right\} - \sum_j \left\{\gamma_j \log(\gamma'_j) - \log\left(\Gamma(\gamma_j)\right) + \left(\gamma_j - 1\right)\left[\psi\left(\gamma_j\right) - \log\left(\gamma'_j\right)\right] - \gamma'_j\left[\frac{\gamma_j}{\gamma'_j}\right]\right\} \\
&+ \sum_i \left\{\alpha\log(\alpha) - \log\left(\Gamma(\alpha)\right) + (\alpha - 1)\left[\psi\left(\zeta_i\right) - \log\left(\zeta'_i\right)\right] - \alpha\left[\frac{\zeta_i}{\zeta'_i}\right]\right\} \\
&- \sum_i \left\{\zeta_i \log(\zeta'_i) - \log\left(\Gamma(\zeta_i)\right) + \left(\zeta_i - 1\right)\left[\psi\left(\zeta_i\right) - \log\left(\zeta'_i\right)\right] - \zeta'_i\left[\frac{\zeta_i}{\zeta'_i}\right]\right\} \\
&+ \sum_{k=1}^{K_{\max}} \sum_{a=1}^{20} \left(\textstyle\sum_i \phi_i^k\right)\omega_{ij}\iota_a^k\left(\psi\left(\lambda_a^k\right) - \psi\left(\sum_{a'=1}^{20} \lambda_{a'}^k\right)\right) + \sum_i \left(\sum_j \omega_{ij} \sum_{k=1}^{\infty} p\left(z_i = k\right)\right)\left(\psi\left(\zeta_i\right) - \log\left(\zeta'_i\right)\right) \\
&+ \sum_j \left(\sum_i \omega_{ij} \sum_{k=1}^{\infty} p\left(z_i = k\right)\right)\left(\psi\left(\gamma_j\right) - \log\left(\gamma'_j\right)\right) - \sum_{ij}\left(\frac{\gamma_j}{\gamma'_j}\right)\left(\frac{\zeta_i}{\zeta'_i}\right)
\end{aligned} \tag{2}$$

### 2 Stochastic Optimization of the Variational Parameters

#### 2.1 Compute the variational function parameters for position of site i to be reallocated to cluster k

Following the principles of the variational inference (Bishop. 2006, Blei and Jordan. 2006, Hoffman 2013), variational parameters of $z_i^k$ is the local parameters, we consider the derivation of the update equation for only one variable by fixing the others' distribution. The optimal coordinate updates are derived for local parameters. The log of the optimized factor to the posterior distribution of $z_i^k$ is

$$\log q^*\left(z\right) = E_{V,\pi,l,r}\left[\log p\left(\Xi, V, z, \pi, l, r\right)\right] + const = \sum_i \sum_{k=1}^{K} z_i^k \log\left(\theta_i^k\right) + const$$

$$\begin{aligned}
\log\theta_i^k &= \mathrm{E}_q\left[\log\left(V_k\right)\right] + \sum_{k'=1}^{k-1} \mathrm{E}_q\left[\log\left(1 - V_{k'}\right)\right] + \sum_{a=1}^{20} \iota_a^k \mathrm{E}_q\left[\log\left(\pi_a^k\right)\right] \\
&= \left\{\psi\left(\vartheta_k\right) - \psi\left(\vartheta_k + \vartheta'_k\right)\right\} + \left\{\sum_{k'=1}^{k-1} \psi\left(\vartheta'_{k'}\right) - \psi\left(\vartheta_{k'} + \vartheta'_{k'}\right)\right\} \\
&+ \sum_{a=1}^{20} \iota_a^k\left\{\psi\left(\lambda_a^k\right) - \psi\left(\sum_{a'=1}^{20} \lambda_{a'}^k\right)\right\}
\end{aligned}$$

$$\mathrm{E}_q\left[z_i^k\right] = \phi_i^k = \exp\left\{\log\left(\theta_i^k\right)\right\}$$

#### 2.2 Updating stick-breaking representation

Similarity, the optimized solution to the posterior distribution of unit length sticks $V_k$

$$\log q^*(V) = E_{z,\pi,l,r}\left[\log p\left(\Xi, z, \pi, l, r\right)\right] + const$$

$$\begin{aligned}&\log q^*(V_k) = (1-1)\log(V_k) + (\kappa - 1)\log(1 - V_k)\\&+\sum_i \mathrm{E}_q\left[z_i^k\right]\log(V_k) + \sum_i \sum_{k'=k+1}^{K_{\max}} E_q\left[z_i^{k'}\right]\log(1-V_k) + const\end{aligned} \tag{3}$$

which has the logarithmic form of the Beta distribution and the corresponding variational distributions of breaking proportions $q(V_k)$ are considered to be the Beta distribution. Then, the standard conditions are satisfied for a closed form coordinate update for local parameters. Based on this framework, the variational parameters of $V_k$ are updated by computing variational expectations in equation (3) (Blei and Jordan. 2006, Hoffman et al. 2013),

$$\vartheta_k = 1 + \sum_i \mathrm{E}_q\left[z_i^k\right] = 1 + \sum_i \phi_i^k$$

$$\vartheta_k' = \kappa + \sum_i \sum_{k'=k+1}^{K_{\max}} \mathrm{E}_q\left[z_i^{k'}\right] = \kappa + \sum_i \sum_{k'=k+1}^{K_{\max}} \phi_i^{k'}$$

#### 2.3 Updating equilibrium frequencies profile

Based on the principal framework of stochastic variational inference (Hoffman et al. 2013), we consider parameters of equilibrium frequencies profile as global parameters. These parameters are updated by a stochastic gradient step and noisy estimations of the natural gradient of the variational objective with respect to $\pi_a^k$. Following the computational method of natural gradient (Hoffman et al. 2013), we compute the natural gradient of equation (2) with respect to the global variational parameters of $\pi_a^k$.
We follow (Lartillot 2004, 2013) to choose Dirichlet distribution as a prior distribution with concentration parameters $\upsilon_a$, so a conditional distribution of profile given the observation data has form of the Dirichlet distribution. We consider the representation of the exponential family,

$$p\left(\pi|\Xi, z, l, r\right) = h(\pi)\exp\left(\eta\left(\Xi, z, l, r\right)^T t(\pi) - a\left(\eta\left(\Xi, z, l, r\right)\right)\right)$$

where: $h(.)$ is the base measure; $a(.)$ is the log-normalize; $\eta(.)$ is the natural parameter; $t(.)$ is the sufficient statistics.
As above assumption of variational parameters, we set $q(\pi|\lambda)$ to be Dirichlet distribution as the complete conditional distributions. So we get

$$q(\pi|\lambda) = h(\pi)\exp\left(\lambda^T t(\pi) - a(\lambda)\right)$$

We consider the lower bound in equation (2) for only profile variable,

$$\begin{aligned}\mathcal{L}(\pi) &= \mathrm{E}_q\left[\log p\left(\pi|\Xi, z, l, r\right)\right] - \mathrm{E}_q\left[q(\pi|\lambda)\right]\\&= \mathrm{E}_q\left[\log\{h(\pi)\} + \eta\left(\Xi, z, l, r\right)^T t(\pi) - a\left(\eta\left(\Xi, z, l, r\right)\right) - \log\{h(\pi)\} - \lambda^T t(\pi) + a(\lambda)\right]\\&= \mathrm{E}_q\left[\eta\left(\Xi, z, l, r\right)^T t(\pi) - a\left(\eta\left(\Xi, z, l, r\right)\right) - \lambda^T t(\pi) + a(\lambda)\right]\\&= \mathrm{E}_q\left[\eta\left(\Xi, z, l, r\right)\right]^T\left[\nabla_\lambda\{a(\lambda)\}\right] - a\left(\eta\left(\Xi, z, l, r\right)\right) - \lambda^T\left[\nabla_\lambda\{a(\lambda)\}\right] + a(\lambda)\end{aligned}$$

where, the expected value of the sufficient statistics is the gradient of log normalizer $\mathrm{E}_q\left[t(\pi)\right] = \nabla_\lambda\{a(\lambda)\}$
If the classical principal of gradient method for maximization is used directly to find a maximum of $\mathcal{L}(\pi)$ based on taking step of size $\rho$ in direction of the gradient, the optimized results is

$$\lambda^{(t+1)} = \lambda^{(t)} + \rho\nabla_\lambda L(\pi) = \lambda^{(t)} + \rho^{(t)}\left[\nabla_\lambda^2\{a(\lambda)\}\right]\left\{E_q\left[\eta\left(\Xi, z, l, r\right)\right]^T - \lambda^T\right\}$$

Following (Hoffman et al. 2013), by premultiplying the gradient by the inverse Fisher information $\mathrm{G}(\lambda)$ and apply the stochastic natural gradient of the variational objective with respect to profile, we get

$$G(\lambda) = E_\lambda\left[\left(\nabla_\lambda \log q(\pi|\lambda)\right)\left(\nabla_\lambda \log q(\pi|\lambda)\right)^T\right] = \nabla_\lambda^2\{a(\lambda)\}$$

$$\widehat{\nabla_\lambda} L(\pi) = \{G(\lambda)\}^{-1}\nabla_\lambda L(\pi) = \left\{E_q\left[\eta\left(\Xi, z, l, r\right)\right] - \lambda\right\}$$

$$\begin{aligned}\lambda^{(t+1)} &= \lambda^{(t)} + \rho\widehat{\nabla_\lambda} L(\pi) = \lambda^{(t)} + \rho\left\{E_q\left[\eta(\Xi, z, l, r)\right] - \lambda^{(t)}\right\} \\ &= (1-\rho)\lambda^{(t)} + \rho\left\{E_q\left[\eta(\Xi, z, l, r)\right]\right\}\end{aligned} \tag{4}$$

Based traditional variational inference, we get the conditional distribution of profile

$$\log p(\pi|z, l, r, \Xi) = \log\left(\pi_a^k\right) \times \sum_i \mathrm{E}_q\left[z_i^k\right] \times \iota_a^k + \log\left(\pi_a^k\right)(\upsilon_a - 1) + const$$

$$\mathrm{E}_q\left[\eta(\Xi, z, l, r)\right] = \upsilon_a + \sum_i \mathrm{E}_q\left[z_i^k\right] \times \iota_a^k \tag{5}$$

By substituting equation (5) into equation (4), the final result of updated variational parameters of site-specific profile is

$$\left(\lambda_a^k\right)^{(t+1)} = \left(1 - \rho^{(t)}\right)\left(\lambda_a^k\right)^{(t)} + \rho^{(t)}\left\{\nu_a + \sum_i \phi_i^k \times \iota_a^k\right\}$$

#### 2.4 Updating branch lengths

Similarity, the conditional distribution given the observational data is

$$\mathrm{E}_{q(r),q(\Xi)}\left[\,\log p(l|\Xi, z, \pi, r)\,\right] = \sum_j \left(1 - 1 + \sum_i \omega_{ij}\right)\log(l_j) - \left(\beta - \sum_i \mathrm{E}_q[r_i]\right) l_j + const$$

$$E_q\left[\eta(\Xi, z, \pi, r)\right] = \begin{bmatrix} 1 - 1 + \sum\limits_i \omega_{ij} \\ \beta + \sum\limits_i \mathrm{E}_q[r_i] \end{bmatrix}$$

The stochastic natural gradient of the variational objective with respect to the branch length

$$\begin{aligned}(\gamma_j)^{(t+1)} &= \left(1 - \rho^{(t)}\right)(\gamma_j)^{(t)} + \rho^{(t)}\left\{1 + \sum_i \omega_{ij}\right\} \\ \left(\gamma_j^{'}\right)^{(t+1)} &= \left(1 - \rho^{(t)}\right)\left(\gamma_j^{'}\right)^{(t)} + \rho^{(t)}\left\{\beta + \sum_i \frac{\zeta_i}{\zeta_i^{'}}\right\}\end{aligned}$$

#### 2.5 Updating rate across site

Similarity, the conditional distribution given the observational data is

$$\mathrm{E}_{q(l),q(\Xi)}\left[\,\log p(r|\Xi, z, \pi, l)\,\right] = \sum_i \left(\alpha - 1 + \sum_j \omega_{ij}\right)\log(r_i) - \left(\alpha - \sum_j \mathrm{E}_q[l_j]\right) r_i + const$$

$$\mathrm{E}_q\left[\eta(\Xi, z, \pi, l)\right] = \begin{bmatrix} \alpha - 1 + \sum\limits_j \omega_{ij} \\ \alpha + \sum\limits_j \mathrm{E}_q[l_j] \end{bmatrix}$$

The stochastic natural gradient of the variational objective with respect to the rate across sites

$$\begin{aligned}(\zeta_i)^{(t+1)} &= \left(1 - \rho^{(t)}\right)(\zeta_i)^{(t)} + \rho^{(t)}\left\{\alpha + \sum_j \omega_{ij}\right\} \\ \left(\zeta_i^{'}\right)^{(t+1)} &= \left(1 - \rho^{(t)}\right)\left(\zeta_i^{'}\right)^{(t)} + \rho^{(t)}\left\{\alpha + \sum_j \frac{\gamma_j}{\gamma_j^{'}}\right\}\end{aligned}$$

#### 2.6 Compute the variational function parameters for the number of substitution events

During a time interval, the number of substitution events is distributed according to a Poisson distribution of rate $r_i l_j$ (Lartillot 2006). This is a prior distribution for $n_{ij}$:

$$p(n|r, l) = \prod_i \prod_j e^{(-r_i l_j)} \frac{(r_i l_j)^{n_{ij}}}{n_{ij}!}$$

The posterior distribution of $n_{ij}$ during a time interval, given that the initial and final states ($\sigma_i = a$ and $\sigma_f = b$) at the process at time evolution $r_i l_j$, is as follow:

$$p\left(n|a,b,\pi,r,l\right) = \frac{p\left(b|a,\pi,n\right)p\left(n|r,l\right)}{\sum\limits_{n\geq 0} p\left(b|a,\pi,n\right)p\left(n|r,l\right)}$$

We consider two cases which are proposed (Lartillot 2006):

- If $a \neq b$ a truncated Poisson distribution is considered for $n_{ij}$
- If a = b, the posterior distribution of $n_{ij}$ is computed as follow:

$$\begin{array}{l} p_{a\to b}\left(r_i l_j\right) = p\left(a,b|\pi,r,l\right) = e^{-r_i l_j} + \left(1 - e^{-r_i l_j}\right)\pi_b \\ p\left(n>0|a,b,\pi,r,l\right) = \prod\limits_i \prod\limits_j \frac{\pi_b}{p(a,b|\pi,r_i,l_j)} \frac{e^{-r_i l_j}\left(r_i l_j\right)^{n_{ij}}}{n_{ij}!} \\ p\left(n=0|a,b,\pi,r,l\right) = \prod\limits_i \prod\limits_j \frac{e^{-r_i l_j}}{p(a,b|\pi,r_i,l_j)} \end{array}$$

The log of the optimized factor to the posterior distribution of having n substitutions along a branch of length $l_j$ is

$$\log q^*(n) = \mathrm{E}_{q(\pi),q(z),q(r),q(l)}\left[\log p\left(a,b,\pi,z,r,l,n\right)\right] + const$$

$$\begin{array}{l} \log q^*(n) = \mathrm{E}_{q(z)}(p(z))\mathrm{E}_{q(\pi)}\left[\log p(\pi)\right] + \mathrm{E}_{q(r),q(l)}\left[\log p\left(n|r,l\right)\right] \\ +\mathrm{E}_{q(\pi),q(r),q(l)}\left[\log\left(p\left(a,b|\pi,r,l\right)\right)\right]\mathrm{E}_{q(z)}(p(z)) + const \end{array} \tag{6}$$

The variational parameters of n are updated by computing variational expectation in equation (6) as follow:

$$\begin{array}{l} \omega_{ij} = \sum\limits_{k=1}^{K_{max}} \sum\limits_{a=1}^{20} \phi_i^k \left[\psi\left(\lambda_a^k\right) - \psi\left(\sum\limits_{a'=1}^{20} \lambda_{a'=1}^k\right)\right] \\ + \left[\psi\left(\zeta_i\right) - \log\left(\zeta_i^{'}\right)\right] + \left[\psi\left(\gamma_j\right) - \log\left(\gamma_j^{'}\right)\right] - \left[\frac{\gamma_j}{\gamma_j^{'}} \frac{\zeta_i}{\zeta_i^{'}}\right] \\ + \sum\limits_{k=1}^{K_{max}} \phi_i^k \mathrm{E}_{q(\pi),q(r),q(l)}\left[\log\left(p\left(a,b|\pi,r,l\right)\right)\right] \end{array} \tag{7}$$

#### 2.7 Compute the variational function parameters for the number of amino acid types in each category

Similarity, the optimized solution to the posterior distribution for the number of amino acid types, conditional on the states at the ends is computed as follows:

$$\begin{array}{l} \log q^*\left(w_a^k\right) = \sum\limits_i \mathrm{E}_{q(z)}\left[p(z_i^k)\right]\mathrm{E}_{q(\pi)}\left[\log\left(\pi_a^k\right)\right] + \sum\limits_i \sum\limits_j \mathrm{E}_{q(r),q(l)} \log\left[1 - e^{-r_i l_j}\right] \\ + \sum\limits_i \sum\limits_j \mathrm{E}_{q(z)}\left[p(z_i^k)\right]\mathrm{E}_{q(\pi),q(r),q(l)}\left[\log\left(p\left(a,b|\pi,r_i,l_j\right)\right)\right] \end{array}$$

The variational parameters of $\iota_a^k$ are updated as follows:

$$\begin{array}{l} \iota_a^k = \sum\limits_i \phi_i^k \left[\psi\left(\lambda_a^k\right) - \psi\left(\sum\limits_{a'=1}^{20} \lambda_{a'}^k\right)\right] + \sum\limits_i \sum\limits_j \mathrm{E}_{q(r),q(l)} \log\left[1 - e^{-r_i l_j}\right] \\ + \sum\limits_i \sum\limits_j \phi_i^k \mathrm{E}_{q(\pi),q(r),q(l)}\left[\log\left(p\left(a,b|\pi,r_i,l_j\right)\right)\right] \end{array} \tag{8}$$

The expected logarithm of the functions $\mathrm{E}_{q(\pi),q(r),q(l)}\left[\log\left(p\left(a,b|\pi,r_i,l_j\right)\right)\right]$ and $\mathrm{E}_{q(r),q(l)}\left[\log\left(1 - e^{-r_i l_j}\right)\right]$ in equation (7, 8) have not the closed form. Thus, calculation of these equations are analytically intractable. To use standard form of variational inference (Bishop. 2006) or stochastic variational inference (Hoffman et al. 2013), we need a closed-form expression. In order to overcome these problems, we consider a common approach:

- By applying a first-order Taylor expansion to preserve a bound, intractable expectations are avoided. By using these results, the variational parameters of the number of substitution and the number of type of amino acid are updated as follow:

$$\omega_{ij} \approx \sum_{k=1}^{K_{max}} \sum_{a=1}^{20} \phi_i^k \left[\psi\left(\lambda_a^k\right) - \psi\left(\sum_{a'=1}^{20} \lambda_{a'=1}^k\right)\right]$$
$$+\left[\psi\left(\zeta_i\right) - \log\left(\zeta_i'\right)\right] + \left[\psi\left(\gamma_j\right) - \log\left(\gamma_j'\right)\right] - \left[\frac{\gamma_j}{\gamma_j'}\frac{\zeta_i}{\zeta_i'}\right]$$
$$+\sum_a \log\left[e^{-\frac{1}{\beta}} + \left(1 - e^{-\frac{1}{\beta}}\right)\pi_a'\right] + \sum_a \frac{\pi_a' - 1}{\pi_a' e^{\frac{1}{\beta}} - \pi_a' + 1}\left(\frac{\zeta_i}{\zeta_i'}\frac{\gamma_j}{\gamma_j'} - \frac{1}{\beta}\right)$$
$$+\sum_{k=1}^{K_{max}} \sum_{a=1}^{20} \phi_i^k \left(\frac{e^{\frac{1}{\beta}} - 1}{\left(e^{\frac{1}{\beta}} - 1\right)\pi_a' + 1}\right)\left(\left[\psi\left(\lambda_a^k\right) - \psi\left(\sum_{a'=1}^{20} \lambda_{a'=1}^k\right)\right] - \pi_a'\right)$$

$$\iota_a^k \approx \sum_i \phi_i^k \left[\psi\left(\lambda_a^k\right) - \psi\left(\sum_{a'=1}^{20} \lambda_{a'}^k\right)\right]$$
$$+\log\left(1 - e^{-\frac{1}{\beta}}\right) + \left(\frac{1}{e^{\frac{1}{\beta}} - 1}\right)\sum_i \sum_j \left(\frac{\zeta_i}{\zeta_i'}\frac{\gamma_j}{\gamma_j'} - \frac{1}{\beta}\right)$$
$$+\sum_a \log\left[e^{-\frac{1}{\beta}} + \left(1 - e^{-\frac{1}{\beta}}\right)\pi_a'\right] + \sum_a \frac{\pi_a' - 1}{\pi_a' e^{\frac{1}{\beta}} - \pi_a' + 1}\left(\frac{\zeta_i}{\zeta_i'}\frac{\gamma_j}{\gamma_j'} - \frac{1}{\beta}\right)$$
$$+\sum_i \phi_i^k \sum_{a=1}^{20} \left(\frac{e^{\frac{1}{\beta}} - 1}{\left(e^{\frac{1}{\beta}} - 1\right)\pi_a' + 1}\right)\left(\left[\psi\left(\lambda_a^k\right) - \psi\left(\sum_{a'=1}^{20} \lambda_{a'=1}^k\right)\right] - \pi_a'\right)$$

#### B. Variational Inference for JC69 model

The D data are two aligned sequences, each n sites long, with x difference. The likelihood is given by the JC69 model as:

$$p\left(D|d\right) = \left(\frac{1}{4}p_1\right)^x \left(\frac{1}{4}p_0\right)^{n-x} = \left(\frac{1}{16} - \frac{1}{16}e^{\frac{-4d}{3}}\right)^x \left(\frac{1}{16} + \frac{3}{16}e^{\frac{-4d}{3}}\right)^{n-x}$$

In Variational inference framework, the optimal solution to the posterior distribution of d is

$$\log q^*\left(d|\gamma,\lambda\right) = \mathrm{E}_q\left[\log p\left(d\right) + \log\left(D|d\right)\right]$$
$$= \mathrm{E}_q\left[x \log\left(\frac{1}{16} - \frac{1}{16}e^{\frac{-4d}{3}}\right)\right] + \mathrm{E}_q\left[\left(n-x\right)\log\left(\frac{1}{16} + \frac{3}{16}e^{\frac{-4d}{3}}\right)\right]$$
$$+\mathrm{E}_q\left[\log\left(\frac{(\beta)^\alpha}{\Gamma(\alpha)}\right) + (\alpha - 1)\log\left(d\right) - \beta d\right]$$

We consider second-order Taylor expansion for $\log\left(\frac{1}{16} - \frac{1}{16}e^{\frac{-4d}{3}}\right)$ and $\log\left(\frac{1}{16} + \frac{3}{16}e^{\frac{-4d}{3}}\right)$ for d at $d' = \frac{\gamma}{\lambda}$

$$\log\left(\frac{1}{16} - \frac{1}{16}e^{\frac{-4d}{3}}\right) \approx \log\left(\frac{1}{16} - \frac{1}{16}e^{\frac{-4d'}{3}}\right) + \frac{\partial \log\left(\frac{1}{16} - \frac{1}{16}e^{\frac{-4d}{3}}\right)}{\partial d}\left(d - d'\right)$$
$$+\left(d - d'\right)\frac{\partial^2 \log\left(\frac{1}{16} - \frac{1}{16}e^{\frac{-4d}{3}}\right)}{\partial d^2}\left(d - d'\right)$$

$$\frac{\partial \log\left(\frac{1}{16}-\frac{1}{16}e^{\frac{-4d}{3}}\right)}{\partial d}|_{d=d'} = \frac{1}{\frac{1}{16}-\frac{1}{16}e^{\frac{-4d'}{3}}}\frac{\partial\left(\frac{1}{16}-\frac{1}{16}e^{\frac{-4d}{3}}\right)}{\partial d}|_{d=d'}$$

$$= \frac{e^{\frac{-4d'}{3}}}{12\left(\frac{1}{16}-\frac{1}{16}e^{\frac{-4d'}{3}}\right)} = \frac{4}{3e^{\frac{4d'}{3}}-3}$$

$$\frac{\partial^2 \log\left(\frac{1}{16}-\frac{1}{16}e^{\frac{-4d'}{3}}\right)}{\partial d^2}|_{d=d'} = 4\frac{\partial}{\partial d}\left(\frac{1}{3e^{\frac{4d}{3}}-3}\right) = -\frac{16e^{\frac{4d'}{3}}}{\left(3e^{\frac{4d'}{3}}-3\right)^2}$$

$$\log\left(\frac{1}{16}+\frac{3}{16}e^{\frac{-4d}{3}}\right) \approx \log\left(\frac{1}{16}+\frac{3}{16}e^{\frac{-4d'}{3}}\right) + \frac{\partial \log\left(\frac{1}{16}+\frac{3}{16}e^{\frac{-4d}{3}}\right)}{\partial d}\left(d-d'\right)$$
$$+\left(d-d'\right)\frac{\partial^2 \log\left(\frac{1}{16}+\frac{3}{16}e^{\frac{-4d}{3}}\right)}{\partial d^2}\left(d-d'\right)$$

$$\frac{\partial \log\left(\frac{1}{16}+\frac{3}{16}e^{\frac{-4d}{3}}\right)}{\partial d}|_{d=d'} = \frac{1}{\frac{1}{16}+\frac{3}{16}e^{\frac{-4d'}{3}}}\frac{\partial\left(\frac{1}{16}+\frac{3}{16}e^{\frac{-4d}{3}}\right)}{\partial d}|_{d=d'}$$

$$= -\frac{e^{\frac{-4d'}{3}}}{4\left(\frac{1}{16}+\frac{3}{16}e^{\frac{-4d'}{3}}\right)} = -\frac{4}{e^{\frac{4d'}{3}}+3}$$

$$\frac{\partial^2 \log\left(\frac{1}{16}+\frac{3}{16}e^{\frac{-4d}{3}}\right)}{\partial d^2}|_{d=d'} = -4\frac{\partial}{\partial d}\left(\frac{1}{e^{\frac{4d}{3}}+3}\right) = \frac{16e^{\frac{4d'}{3}}}{3\left(e^{\frac{4d'}{3}}+3\right)^2}$$

With result of first-order Taylor expansion and the principle of the VI framework, the optimal solution to the posterior distribution of d is

$$\log q^*(d|\gamma,\lambda) \approx x\mathrm{E}_q\left[\log\left(\frac{1}{16}-\frac{1}{16}e^{\frac{-4d'}{3}}\right)+\left(\frac{4}{3e^{\frac{4d'}{3}}-3}\right)\left(d-d'\right)\right]$$
$$+(n-x)\,\mathrm{E}_q\left[\log\left(\frac{1}{16}+\frac{3}{16}e^{\frac{-4d'}{3}}\right)+\left(-\frac{4}{e^{\frac{4d'}{3}}+3}\right)\left(d-d'\right)\right]$$
$$+\mathrm{E}_q\left[\log\left(\frac{(\beta)^{\alpha}}{\Gamma(\alpha)}\right)+(\alpha-1)\log(d)-\beta d\right]$$

$$\log q^*(d|\gamma,\lambda) \approx x\mathrm{E}_q\left[\left(\frac{4}{3e^{\frac{4d'}{3}}-3}\right)d\right]+(n-x)\,\mathrm{E}_q\left[\left(-\frac{4}{e^{\frac{4d'}{3}}+3}\right)d\right]$$
$$+\mathrm{E}_q\left[(\alpha-1)\log(d)-\beta d\right]+const$$

which has the logarithm form of the gamma distribution.

#### C. Variational Inference for K80 model

Here we illustrate the major features of Variational Inference by applying it to the problem of estimating d and the transition/transversion rate ratio $\kappa$ under the K80 model using a pair of DNA sequences. D is an alignment of the human and orangutan mitochondrial 12S rRNA genes, summarized as $n_S = 84$ transitional differences and $n_V = 6$ transversional differences at n = 948 sites. We assign independent gamma priors,

$$p(d) = Gamma\left(d|\alpha_d,\beta_d\right) = \frac{(\beta_d)^{\alpha_d}}{\Gamma(\alpha_d)}d^{\alpha_d-1}e^{-\beta_d d} with \alpha_d = 2, \beta_d = 20$$
$$p(\kappa) = Gamma\left(\kappa|\alpha_\kappa,\beta_\kappa\right) = \frac{(\beta_\kappa)^{\alpha_\kappa}}{\Gamma(\alpha_\kappa)}\kappa^{\alpha_\kappa-1}e^{-\beta_\kappa \kappa} with \alpha_\kappa = 2, \beta_\kappa = 0.1$$

The likelihood is given by the K80 model as:

$$p\left(D|d,\kappa\right) = \left(\frac{1}{4}p_0\right)^{n-n_S-n_V}\left(\frac{1}{4}p_1\right)^{n_S}\left(\frac{1}{4}p_2\right)^{n_V}$$

$$p_0 = \frac{1}{4}+\frac{1}{4}e^{-\frac{4d}{\kappa+2}}+\frac{1}{2}e^{-2d\frac{\kappa+1}{\kappa+2}}$$
$$p_1 = \frac{1}{4}+\frac{1}{4}e^{-\frac{4d}{\kappa+2}}-\frac{1}{2}e^{-2d\frac{\kappa+1}{\kappa+2}}$$
$$p_2 = \frac{1}{4}-\frac{1}{4}e^{-\frac{4d}{\kappa+2}}$$

In Variational inference framework, the optimal solution to the posterior distribution of d is

$$\log q^*\left(d|\gamma,\gamma'\right) = \mathrm{E}_q\left[\log p\left(d\right)+\log\left(D|d,\kappa\right)\right]$$
$$= \mathrm{E}_q\left[(n-n_S-n_V)\log\left(\frac{1}{16}+\frac{1}{16}e^{-\frac{4d}{\kappa+2}}+\frac{1}{8}e^{-2d\frac{\kappa+1}{\kappa+2}}\right)\right]$$
$$+\mathrm{E}_q\left[(n_S)\log\left(\frac{1}{16}+\frac{1}{16}e^{-\frac{4d}{\kappa+2}}-\frac{1}{8}e^{-2d\frac{\kappa+1}{\kappa+2}}\right)\right]$$
$$+\mathrm{E}_q\left[(n_V)\log\left(\frac{1}{16}-\frac{1}{16}e^{-\frac{4d}{\kappa+2}}\right)\right]$$
$$+\mathrm{E}_q\left[\log\left(\frac{(\beta_d)^{\alpha_d}}{\Gamma(\alpha_d)}\right)+(\alpha_d-1)\log(d)-\beta_d d\right]$$

$$\log q^*\left(\kappa|\lambda,\lambda^{'}\right) = \mathrm{E}_q\left[\log p\left(\kappa\right)+\log\left(D|d,\kappa\right)\right]$$
$$= \mathrm{E}_q\left[\left(n-n_S-n_V\right)\log\left(\frac{1}{16}+\frac{1}{16}e^{-\dfrac{4d}{\kappa+2}}+\frac{1}{8}e^{-2d\dfrac{\kappa+1}{\kappa+2}}\right)\right]$$
$$+\mathrm{E}_q\left[\left(n_S\right)\log\left(\frac{1}{16}+\frac{1}{16}e^{-\dfrac{4d}{\kappa+2}}-\frac{1}{8}e^{-2d\dfrac{\kappa+1}{\kappa+2}}\right)\right]$$
$$+\mathrm{E}_q\left[\left(n_V\right)\log\left(\frac{1}{16}-\frac{1}{16}e^{-\dfrac{4d}{\kappa+2}}\right)\right]$$
$$+\mathrm{E}_q\left[\log\left(\frac{\left(\beta_\kappa\right)^{\alpha_\kappa}}{\Gamma(\alpha_\kappa)}\right)+\left(\alpha_\kappa-1\right)\log\left(\kappa\right)-\beta_\kappa\kappa\right]$$

We consider second-order Taylor expansion for $\log\left(\frac{1}{16}+\frac{1}{16}e^{-\frac{4d}{\kappa+2}}+\frac{1}{8}e^{-2d\frac{\kappa+1}{\kappa+2}}\right)$ and $\log\left(\frac{1}{16}+\frac{1}{16}e^{-\frac{4d}{\kappa+2}}-\frac{1}{8}e^{-2d\frac{\kappa+1}{\kappa+2}}\right)$ and $\log\left(\frac{1}{16}-\frac{1}{16}e^{-\frac{4d}{\kappa+2}}\right)$ for d and $\kappa$ at $d^{'}=\frac{\gamma}{\gamma^{'}}$ and $\kappa^{'}=\frac{\lambda}{\lambda^{'}}$

For $\log\left(\frac{1}{16}+\frac{1}{16}e^{-\frac{4d}{\kappa+2}}+\frac{1}{8}e^{-2d\frac{\kappa+1}{\kappa+2}}\right)$

$$\log\left(\frac{1}{16}+\frac{1}{16}e^{-\dfrac{4d}{\kappa+2}}+\frac{1}{8}e^{-2d\dfrac{\kappa+1}{\kappa+2}}\right)\approx\log\left(\frac{1}{16}+\frac{1}{16}e^{-\dfrac{4d^{'}}{\kappa^{'}+2}}+\frac{1}{8}e^{-2d^{'}\dfrac{\kappa^{'}+1}{\kappa^{'}+2}}\right)$$
$$+\frac{\partial\log\left(\frac{1}{16}+\frac{1}{16}e^{-\dfrac{4d}{\kappa+2}}+\frac{1}{8}e^{-2d\dfrac{\kappa+1}{\kappa+2}}\right)}{\partial d}\left(d-d^{'}\right)+\frac{\partial\log\left(\frac{1}{16}+\frac{1}{16}e^{-\dfrac{4d}{\kappa+2}}+\frac{1}{8}e^{-2d\dfrac{\kappa+1}{\kappa+2}}\right)}{\partial\kappa}\left(\kappa-\kappa^{'}\right)$$
$$+\frac{\partial^2\log\left(\frac{1}{16}+\frac{1}{16}e^{-\dfrac{4d}{\kappa+2}}+\frac{1}{8}e^{-2d\dfrac{\kappa+1}{\kappa+2}}\right)}{\partial d\partial\kappa}\left(d-d^{'}\right)\left(\kappa-\kappa^{'}\right)$$
$$+\frac{1}{2}\left(d-d^{'}\right)\frac{\partial^2\log\left(\frac{1}{16}+\frac{1}{16}e^{-\dfrac{4d}{\kappa+2}}+\frac{1}{8}e^{-2d\dfrac{\kappa+1}{\kappa+2}}\right)}{\partial d^2}\left(d-d^{'}\right)$$
$$+\frac{1}{2}\left(\kappa-\kappa^{'}\right)\frac{\partial^2\log\left(\frac{1}{16}+\frac{1}{16}e^{-\dfrac{4d}{\kappa+2}}+\frac{1}{8}e^{-2d\dfrac{\kappa+1}{\kappa+2}}\right)}{\partial\kappa^2}\left(\kappa-\kappa^{'}\right)$$

$$\frac{\partial \log\left(\frac{1}{16}+\frac{1}{16}e^{-\frac{4d}{\kappa+2}}+\frac{1}{8}e^{-2d\frac{\kappa+1}{\kappa+2}}\right)}{\partial d}\Big|_{d=d',\kappa=\kappa'}$$

$$=\frac{1}{\frac{1}{16}+\frac{1}{16}e^{-\frac{4d'}{\kappa'+2}}+\frac{1}{8}e^{-2d'\frac{\kappa'+1}{\kappa'+2}}}\frac{\partial\left(\frac{1}{16}+\frac{1}{16}e^{-\frac{4d}{\kappa+2}}+\frac{1}{8}e^{-2d\frac{\kappa+1}{\kappa+2}}\right)}{\partial d}\Big|_{d=d',\kappa=\kappa'}$$

$$=\frac{1}{\frac{1}{16}+\frac{1}{16}e^{-\frac{4d'}{\kappa'+2}}+\frac{1}{8}e^{-2d'\frac{\kappa'+1}{\kappa'+2}}}\left(-\frac{\left(\kappa'+1\right)e^{-2d'\frac{\kappa'+1}{\kappa'+2}}}{4\left(\kappa'+2\right)}-\frac{e^{-\frac{4d'}{\kappa'+2}}}{4\left(\kappa'+2\right)}\right)$$

$$=-\frac{4\left(e^{\frac{2d'\left(\kappa'+1\right)}{\kappa'+2}}+\left(\kappa'+1\right)e^{\frac{4d'}{\left(\kappa'+2\right)}}\right)}{\left(\kappa'+2\right)\left(\left(e^{\frac{4d'}{\left(\kappa'+2\right)}}+1\right)e^{\frac{2d'\left(\kappa'+1\right)}{\kappa'+2}}+2e^{\frac{4d'}{\left(\kappa'+2\right)}}\right)}$$

$$\frac{\partial \log\left(\frac{1}{16}+\frac{1}{16}e^{-\frac{4d}{\kappa+2}}+\frac{1}{8}e^{-2d\frac{\kappa+1}{\kappa+2}}\right)}{\partial \kappa}\Big|_{d=d',\kappa=\kappa'}$$

$$=\frac{1}{\frac{1}{16}+\frac{1}{16}e^{-\frac{4d'}{\kappa'+2}}+\frac{1}{8}e^{-2d'\frac{\kappa'+1}{\kappa'+2}}}\frac{\partial\left(\frac{1}{16}+\frac{1}{16}e^{-\frac{4d}{\kappa+2}}+\frac{1}{8}e^{-2d\frac{\kappa+1}{\kappa+2}}\right)}{\partial \kappa}\Big|_{d=d',\kappa=\kappa'}$$

$$=\frac{1}{\frac{1}{16}+\frac{1}{16}e^{-\frac{4d'}{\kappa'+2}}+\frac{1}{8}e^{-2d'\frac{\kappa'+1}{\kappa'+2}}}\left(\frac{d'e^{-\frac{4d'}{\left(\kappa'+2\right)}}}{4\left(\kappa'+2\right)^2}-\frac{d'e^{-\frac{2d'\left(\kappa'+1\right)}{\kappa'+2}}}{4\left(\kappa'+2\right)^2}\right)$$

$$=\frac{4d'\left(e^{\frac{2d'\left(\kappa'+1\right)}{\kappa'+2}}-e^{\frac{4d'}{\kappa'+2}}\right)}{\left(\kappa'+2\right)^2\left(\left(e^{\frac{4d'}{\left(\kappa'+2\right)}}+1\right)e^{\frac{2d'\left(\kappa'+1\right)}{\kappa'+2}}+2e^{\frac{4d'}{\left(\kappa'+2\right)}}\right)}$$

For $\log\left(\frac{1}{16}+\frac{1}{16}e^{-\frac{4d}{\kappa+2}}-\frac{1}{8}e^{-2d\frac{\kappa+1}{\kappa+2}}\right)$

$$\log\left(\frac{1}{16}+\frac{1}{16}e^{-\dfrac{4d}{\kappa+2}}-\frac{1}{8}e^{-2d\dfrac{\kappa+1}{\kappa+2}}\right)\approx\log\left(\frac{1}{16}+\frac{1}{16}e^{-\dfrac{4d'}{\kappa'+2}}-\frac{1}{8}e^{-2d'\dfrac{\kappa'+1}{\kappa'+2}}\right)$$

$$+\frac{\partial\log\left(\frac{1}{16}+\frac{1}{16}e^{-\dfrac{4d}{\kappa+2}}-\frac{1}{8}e^{-2d\dfrac{\kappa+1}{\kappa+2}}\right)}{\partial d}\left(d-d'\right)+\frac{\partial\log\left(\frac{1}{16}+\frac{1}{16}e^{-\dfrac{4d}{\kappa+2}}-\frac{1}{8}e^{-2d\dfrac{\kappa+1}{\kappa+2}}\right)}{\partial\kappa}\left(\kappa-\kappa'\right)$$

$$+\frac{\partial^2\log\left(\frac{1}{16}+\frac{1}{16}e^{-\dfrac{4d}{\kappa+2}}-\frac{1}{8}e^{-2d\dfrac{\kappa+1}{\kappa+2}}\right)}{\partial d\partial\kappa}\left(d-d'\right)\left(\kappa-\kappa'\right)$$

$$+\frac{1}{2}\left(d-d'\right)\frac{\partial^2\log\left(\frac{1}{16}+\frac{1}{16}e^{-\dfrac{4d}{\kappa+2}}-\frac{1}{8}e^{-2d\dfrac{\kappa+1}{\kappa+2}}\right)}{\partial d^2}\left(d-d'\right)$$

$$+\frac{1}{2}\left(\kappa-\kappa'\right)\frac{\partial^2\log\left(\frac{1}{16}+\frac{1}{16}e^{-\dfrac{4d}{\kappa+2}}-\frac{1}{8}e^{-2d\dfrac{\kappa+1}{\kappa+2}}\right)}{\partial\kappa^2}\left(\kappa-\kappa'\right)$$

$$\frac{\partial\log\left(\frac{1}{16}+\frac{1}{16}e^{-\dfrac{4d}{\kappa+2}}-\frac{1}{8}e^{-2d\dfrac{\kappa+1}{\kappa+2}}\right)}{\partial d}\Big|_{d=d',\kappa=\kappa'}$$

$$=\frac{1}{\frac{1}{16}+\frac{1}{16}e^{-\dfrac{4d'}{\kappa'+2}}-\frac{1}{8}e^{-2d'\dfrac{\kappa'+1}{\kappa'+2}}}\frac{\partial\left(\frac{1}{16}+\frac{1}{16}e^{-\dfrac{4d}{\kappa+2}}-\frac{1}{8}e^{-2d\dfrac{\kappa+1}{\kappa+2}}\right)}{\partial d}\Big|_{d=d',\kappa=\kappa'}$$

$$=\frac{1}{\frac{1}{16}+\frac{1}{16}e^{-\dfrac{4d'}{\kappa'+2}}-\frac{1}{8}e^{-2d'\dfrac{\kappa'+1}{\kappa'+2}}}\left(\frac{\left(\kappa'+1\right)e^{-2d'\dfrac{\kappa'+1}{\kappa'+2}}}{4\left(\kappa'+2\right)}-\frac{e^{-\dfrac{4d'}{\kappa'+2}}}{4\left(\kappa'+2\right)}\right)$$

$$=\frac{4\left(e^{\dfrac{2d'\left(\kappa'+1\right)}{\kappa'+2}}+\left(-\kappa'-1\right)e^{\dfrac{4d'}{(\kappa'+2)}}\right)}{(\kappa'+2)\left(\left(e^{\dfrac{4d'}{(\kappa'+2)}}+1\right)e^{\dfrac{2d'\left(\kappa'+1\right)}{\kappa'+2}}+2e^{\dfrac{4d'}{(\kappa'+2)}}\right)}$$

$$\frac{\partial \log\left(\frac{1}{16}+\frac{1}{16}e^{-\frac{4d}{\kappa+2}}-\frac{1}{8}e^{-2d\frac{\kappa+1}{\kappa+2}}\right)}{\partial \kappa}\Big|_{d=d',\kappa=\kappa'}$$

$$=\frac{1}{\frac{1}{16}+\frac{1}{16}e^{-\frac{4d'}{\kappa'+2}}-\frac{1}{8}e^{-2d'\frac{\kappa'+1}{\kappa'+2}}}\frac{\partial\left(\frac{1}{16}+\frac{1}{16}e^{-\frac{4d}{\kappa+2}}-\frac{1}{8}e^{-2d\frac{\kappa+1}{\kappa+2}}\right)}{\partial \kappa}\Big|_{d=d',\kappa=\kappa'}$$

$$=\frac{1}{\frac{1}{16}+\frac{1}{16}e^{-\frac{4d'}{\kappa'+2}}-\frac{1}{8}e^{-2d'\frac{\kappa'+1}{\kappa'+2}}}\left(\frac{d'e^{-\frac{4d'}{(\kappa'+2)}}}{4\left(\kappa'+2\right)^2}+\frac{d'e^{-\frac{2d'\left(\kappa'+1\right)}{\kappa'+2}}}{4\left(\kappa'+2\right)^2}\right)$$

$$=\frac{4d'\left(e^{\frac{2d'\left(\kappa'+1\right)}{\kappa'+2}}+e^{\frac{4d'}{\kappa'+2}}\right)}{\left(\kappa'+2\right)^2\left(\left(e^{\frac{4d'}{(\kappa'+2)}}+1\right)e^{\frac{2d'\left(\kappa'+1\right)}{\kappa'+2}}-2e^{\frac{4d'}{(\kappa'+2)}}\right)}$$

For $\log\left(\frac{1}{16}-\frac{1}{16}e^{-\frac{4d}{\kappa+2}}\right)$

$$\log\left(\frac{1}{16}-\frac{1}{16}e^{-\frac{4d}{\kappa+2}}\right)\approx\log\left(\frac{1}{16}-\frac{1}{16}e^{-\frac{4d'}{\kappa'+2}}\right)$$
$$+\frac{\partial\log\left(\frac{1}{16}-\frac{1}{16}e^{-\frac{4d}{\kappa+2}}\right)}{\partial d}\left(d-d'\right)+\frac{\partial\log\left(\frac{1}{16}-\frac{1}{16}e^{-\frac{4d}{\kappa+2}}\right)}{\partial \kappa}\left(\kappa-\kappa'\right)$$
$$+\frac{\partial^2\log\left(\frac{1}{16}-\frac{1}{16}e^{-\frac{4d}{\kappa+2}}\right)}{\partial d\partial\kappa}\left(d-d'\right)\left(\kappa-\kappa'\right)$$
$$+\frac{1}{2}\left(d-d'\right)\frac{\partial^2\log\left(\frac{1}{16}-\frac{1}{16}e^{-\frac{4d}{\kappa+2}}\right)}{\partial d^2}\left(d-d'\right)$$
$$+\frac{1}{2}\left(\kappa-\kappa'\right)\frac{\partial^2\log\left(\frac{1}{16}-\frac{1}{16}e^{-\frac{4d}{\kappa+2}}\right)}{\partial \kappa^2}\left(\kappa-\kappa'\right)$$

$$\frac{\partial \log\left(\frac{1}{16}-\frac{1}{16}e^{-\frac{4d}{\kappa+2}}\right)}{\partial d}|_{d=d',\kappa=\kappa'}$$

$$=\frac{1}{\frac{1}{16}-\frac{1}{16}e^{-\frac{4d'}{\kappa'+2}}}\frac{\partial\left(\frac{1}{16}-\frac{1}{16}e^{-\frac{4d}{\kappa+2}}\right)}{\partial d}|_{d=d',\kappa=\kappa'}$$

$$=\frac{1}{\frac{1}{16}-\frac{1}{16}e^{-\frac{4d'}{\kappa'+2}}}\left(\frac{e^{-\frac{4d'}{\kappa'+2}}}{4\left(\kappa'+2\right)}\right)=\frac{4}{(\kappa'+2)\left(e^{\frac{4d'}{(\kappa'+2)}}-1\right)}$$

$$\frac{\partial \log\left(\frac{1}{16}-\frac{1}{16}e^{-\frac{4d}{\kappa+2}}\right)}{\partial \kappa}|_{d=d',\kappa=\kappa'}$$

$$=\frac{1}{\frac{1}{16}-\frac{1}{16}e^{-\frac{4d'}{\kappa'+2}}}\frac{\partial\left(\frac{1}{16}-\frac{1}{16}e^{-\frac{4d}{\kappa+2}}\right)}{\partial \kappa}|_{d=d',\kappa=\kappa'}$$

$$=-\frac{1}{\frac{1}{16}-\frac{1}{16}e^{-\frac{4d'}{\kappa'+2}}}\left(\frac{d'e^{-\frac{4d'}{\kappa'+2}}}{4\left(\kappa'+2\right)^2}\right)=-\frac{4d'}{(\kappa'+2)^2\left(e^{\frac{4d'}{(\kappa'+2)}}-1\right)}$$

$$\frac{\partial^2 \log\left(\frac{1}{16}-\frac{1}{16}e^{-\frac{4d}{\kappa+2}}\right)}{\partial d^2}$$

$$=\frac{\partial}{\partial d}\left(\frac{4}{(\kappa'+2)\left(e^{\frac{4d}{(\kappa'+2)}}-1\right)}\right)|_{d=d',\kappa=\kappa'}=-\frac{16e^{\frac{4d'}{(\kappa'+2)}}}{(\kappa'+2)^2\left(e^{\frac{4d'}{(\kappa'+2)}}-1\right)^2}$$

$$\frac{\partial^2 \log\left(\frac{1}{16}-\frac{1}{16}e^{-\frac{4d}{\kappa+2}}\right)}{\partial\kappa^2}|_{d=d',\kappa=\kappa'}$$

$$=\frac{\partial}{\partial\kappa}\left(-\frac{4d'}{(\kappa+2)^2\left(e^{\frac{4d'}{(\kappa+2)}}-1\right)}\right)|_{d=d',\kappa=\kappa'}$$

$$=4d'\frac{1}{\left((\kappa'+2)^2\left(e^{\frac{4d'}{(\kappa'+2)}}-1\right)\right)^2}\frac{\partial}{\partial\kappa}\left((\kappa+2)^2\left(e^{\frac{4d'}{(\kappa+2)}}-1\right)\right)|_{d=d',\kappa=\kappa'}$$

$$=4d'\frac{1}{\left((\kappa'+2)^2\left(e^{\frac{4d'}{(\kappa'+2)}}-1\right)\right)^2}\left(2\left(\kappa'+2\right)\left(e^{\frac{4d'}{(\kappa'+2)}}-1\right)-4d'e^{\frac{4d'}{(\kappa'+2)}}\right)$$

$$=\frac{8d'\left(\left(\kappa'-2d'+2\right)e^{\frac{4d'}{(\kappa'+2)}}-\kappa'-2\right)}{(\kappa'+2)^4\left(e^{\frac{4d'}{(\kappa'+2)}}-1\right)^2}$$

$$\frac{\partial^2 \log\left(\frac{1}{16}-\frac{1}{16}e^{-\frac{4d}{\kappa+2}}\right)}{\partial d\partial\kappa}|_{d=d',\kappa=\kappa'}$$

$$=\frac{\partial}{\partial\kappa}\left(\frac{4}{(\kappa+2)\left(e^{\frac{4d'}{(\kappa+2)}}-1\right)}\right)|_{d=d',\kappa=\kappa'}$$

$$=-\frac{4}{\left((\kappa'+2)\left(e^{\frac{4d'}{(\kappa'+2)}}-1\right)\right)^2}\frac{\partial}{\partial\kappa}\left((\kappa+2)\left(e^{\frac{4d'}{(\kappa+2)}}-1\right)\right)|_{d=d',\kappa=\kappa'}$$

$$=-\frac{4}{\left((\kappa'+2)\left(e^{\frac{4d'}{(\kappa'+2)}}-1\right)\right)^2}\left(-\frac{4d'e^{\frac{4d'}{(\kappa'+2)}}}{\kappa'+2}+e^{\frac{4d'}{(\kappa'+2)}}-1\right)$$

$$=-\frac{4\left(\left(\kappa'-4d'+2\right)e^{\frac{4d'}{(\kappa'+2)}}-\kappa'-2\right)}{(\kappa'+2)^3\left(e^{\frac{4d'}{(\kappa'+2)}}-1\right)^2}$$

$$\frac{\partial^2 \log\left(\frac{1}{16}-\frac{1}{16}e^{-\frac{4d}{\kappa+2}}\right)}{\partial d\partial\kappa}|_{d=d',\kappa=\kappa'}$$

$$=\frac{\partial}{\partial d}\left(-\frac{4d}{(\kappa'+2)^2\left(e^{\frac{4d}{(\kappa'+2)}}-1\right)}\right)=-\frac{4}{(\kappa'+2)^2}\frac{\partial}{\partial d}\left(\frac{d}{e^{\frac{4d}{(\kappa'+2)}}-1}\right)|_{d=d',\kappa=\kappa'}$$

$$=\frac{4}{(\kappa'+2)^2\left(e^{\frac{4d}{(\kappa'+2)}}-1\right)^2}\left(\frac{4d'e^{\frac{4d'}{(\kappa'+2)}}}{\kappa'+2}-e^{\frac{4d'}{(\kappa'+2)}}+1\right)$$

$$=\frac{4\left(\left(4d'-\kappa'-2\right)e^{\frac{4d'}{(\kappa'+2)}}+\kappa'+2\right)}{(\kappa'+2)^3\left(e^{\frac{4d'}{(\kappa'+2)}}-1\right)^2}$$

### 3 Reference


1. Bishop. C. M. 2006. Pattern Recognition and Machine Learning. Springer, 1 edition.

2. Felsenstein J. 1981. Evolutionary trees from DNA sequences: a maximum likelihood approach. J. Mol. Evol. 17:368–376.

3. Blei, D and Jordan, M. 2006. Variational inference for Dirichlet process mixtures. Journal of Bayesian Analysis, 1:121–144.

4. Gopalan, P., Hao, W., Blei, D., Storey, J. 2016. Scaling probabilistic models of genetic variation to millions of humans. Nat Genet. 48, 1587-1590.

5. Hoffman, M., Blei, D., Wang, C. & Paisley, J. 2013. Stochastic variational inference. J. Mach. Learn. Res. 14, 1303–1347.

6. Jordan, M., Ghahramani, Z., Jaakkola, T. & Saul, L. 1999. Introduction to variational methods for graphical models. Mach. Learn. 37, 183–233.

7. Lartillot N., Philippe H. 2004. A Bayesian mixture model for across-site heterogeneities in the amino-acid replacement process. Mol. Biol. Evol. 21:1095–1109.

8. Lartillot N. 2006. Conjugate Gibbs sampling for Bayesian phylogenetic models. J. Comput. Biol. 13:1701–1722.

9. Lartillot, N., Rodrigue, N., Stubbs, D. & Richer, J. 2013. PhyloBayes MPI: phylogenetic reconstruction with infinite mixtures of profiles in a parallel environment. Syst. Biol.. 62, 611–615.

10. Nielsen R. 2002. Mapping mutations on phylogenies. Syst. Biol. 51: 729–739.

11. Quang L.S., Gascuel O., Lartillot N. 2008. Empirical profile mixture models for phylogenetic reconstruction. Bioinformatics 24: 2317–2323.

12. Robbins, H. and Monro, S. 1951. A stochastic approximation method. The Annals of Mathematical Statistics 22, 400–407.

13. Wainwright, M. & Jordan, M. 2008. Graphical models, exponential families, and variational inference. Found. Trends Mach. Learn. 1, 1–305.
